# Encoding of melody in the human auditory cortex

**DOI:** 10.1101/2023.10.17.562771

**Authors:** Narayan Sankaran, Matthew K. Leonard, Frederic Theunissen, Edward F. Chang

## Abstract

Melody is a core component of music in which discrete pitches are serially arranged to convey emotion and meaning. Perception of melody varies along several pitch-based dimensions: (1) the absolute pitch of notes, (2) the difference in pitch between successive notes, and (3) the higher-order statistical expectation of each note conditioned on its prior context. While humans readily perceive melody, how these dimensions are collectively represented in the brain and whether their encoding is specialized for music remains unknown. Here, we recorded high-density neurophysiological activity directly from the surface of human auditory cortex while Western participants listened to Western musical phrases. Pitch, pitch-change, and expectation were selectively encoded at different cortical sites, indicating a spatial code for representing distinct dimensions of melody. The same participants listened to spoken English, and we compared evoked responses to music and speech. Cortical sites selective for music were systematically driven by the encoding of expectation. In contrast, sites that encoded pitch and pitch-change used the same neural code to represent equivalent properties of speech. These findings reveal the multidimensional nature of melody encoding, consisting of both music-specific and domain-general sound representations in auditory cortex.

**Teaser:** The human brain contains both general-purpose and music-specific neural populations for processing distinct attributes of melody.

## Introduction

Musical communication is a hallmark of human behavior. To appreciate music, listeners must extract multiple features across varying timescales from a dynamic acoustic signal. This process leverages core spectrotemporal organizing principles of the human auditory system (*1–3*) and may share evolutionary roots with language (*4–6*).

While music varies in its physical features across cultures and genres, a defining feature across popular idioms is the serial arrangement of discrete pitch-units – or notes – to produce the emergent percept of melody (Fig. 1a). Considered in isolation, each note has a *pitch*, perceived along a low-to-high continuum according to its fundamental frequency. Considered within a melodic context, each note is imbued with higher-order attributes, reflecting the integration of prior information at progressively longer timescales (*4, 7–9*). For instance, the magnitude and direction of *pitch-change* between adjacent notes define the melodic interval and contour respectively, linking the isolated sensory attribute of pitch with our perceptual experience of melody (*10–12*). Listeners are also familiar with melody’s statistical structure and use this knowledge to generate *expectations* about the likelihood of upcoming notes conditioned on the prior sequence. In figure 1a for example, the third-to-last note is relatively unexpected, violating the pattern and tonality established earlier in the phrase. The continuum along which melody violates or fulfills our expectations plays a central role in our aesthetic experience of music (*13–18*), and composers will intentionally modulate expectations to systematically generate patterns of tension and resolution as we listen.

**Figure 1.**
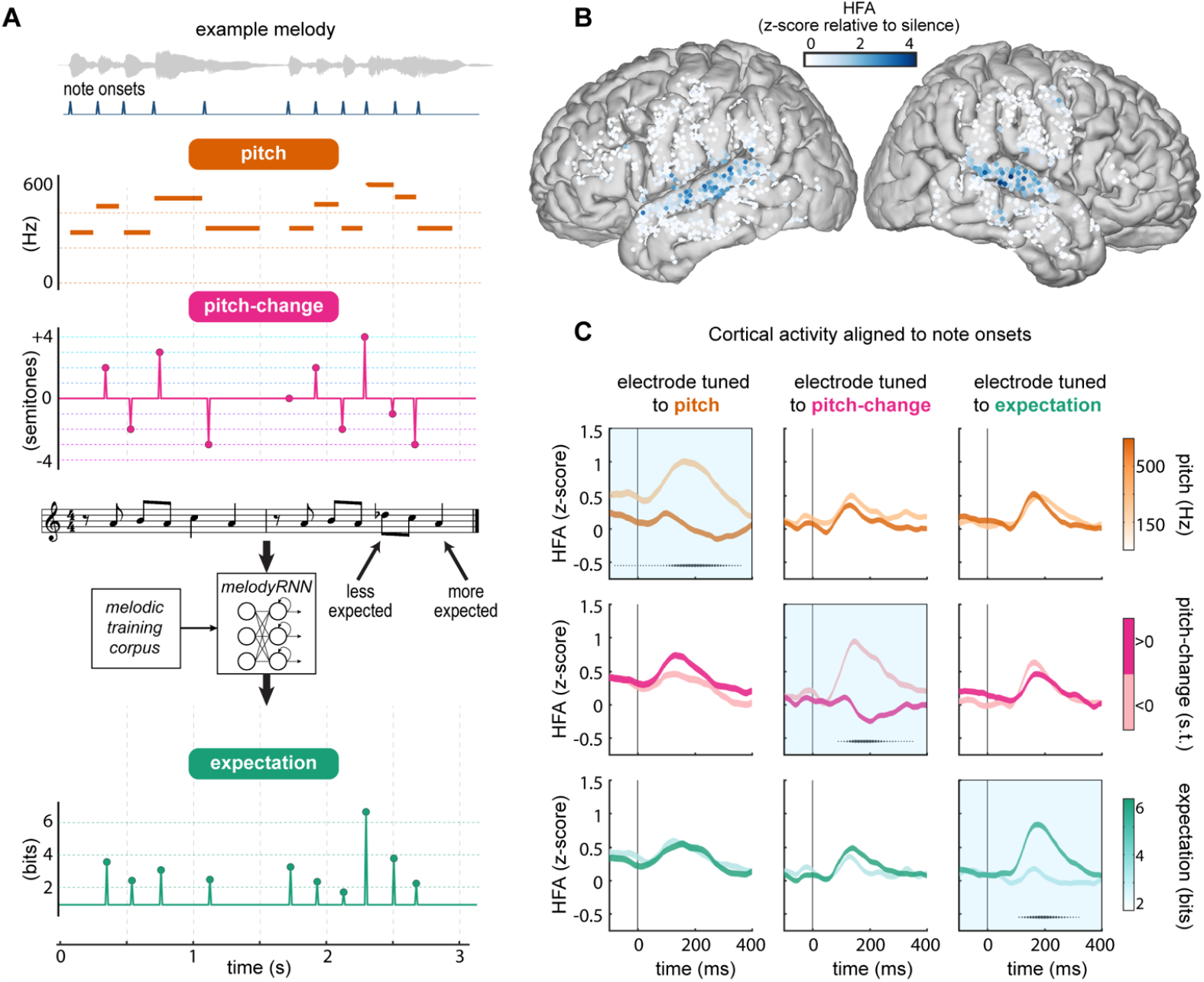
Melodic pitch, pitch-change, and expectation modulate *STG* activity during music listening. **(A)** Three melodic features visualized for an example melody: [1] Absolute pitch (measured in Hz), defined by the fundamental frequency (F0) of each discrete note in the melody. [2] Pitch-change between adjacent notes. The magnitude of change (interval) is measured in semitones, while the direction of change (contour) is binary. [3] Melodic expectation (measured in bits) indicates the surprisal (negative log2 likelihood) of each note, conditioned on preceding notes. Expectation was measured using a pre-trained model of Western melody (melodyRNN). **(B)** Electrodes across all participants (N=8) plotted on a common brain. Color indicates the peak of the average evoked high-frequency activity (HFA) across all phrases. **(C)** Responses at three example electrodes demonstrating distinct tuning to pitch (left column), pitch-change (middle column), and expectation (right column). Each feature distribution is divided into two equal bins (median split). Traces indicate the mean +/- standard error of cortical responses within each bin. Black markers underneath traces indicate timepoints during which responses in the two bins significantly differed (p<0.001; independent two-sample t-test, Bonferroni corrected; marker size indicates t-statistic magnitude).

Although these melodic features – pitch, pitch-change, and expectation – convey distinct perceptual qualities, they stem from the same pitch information computed over progressively longer temporal windows. This raises a fundamental question about whether their representation in the brain is spatially dissociable. Specifically, are different features of melody selectively encoded by distinct neural populations (*19–21*), or do the same populations jointly encode melodic features by modulating afferent pitch input (*22, 23*)? This question has major implications for understanding how the brain represents information at different timescales within dynamic input streams such as music or speech (*24*).

Additionally, the extent to which music is encoded by general auditory mechanisms versus ones specialized for music remains debated (*25–30*). Recent work has found subregions in the human superior temporal gyrus (STG) that selectively respond to music over other sounds like speech (*31–33*). While this is thought to reflect a specialized neural pathway for “music”, it remains unclear which musical properties drive this selectivity. Resolving this question is critical for understanding the nature and extent of specialization in the human brain for important domains of sound.

To address these questions, we used high-density electrocorticography (ECoG) to record neural activity on the human cortical surface while participants listened to a set of Western monophonic musical phrases (see methods for stimulus details). These direct high-density recordings are necessary to resolve fine-grained spatial tuning over millimeters of cortex to dynamic information changing over milliseconds. Within auditory cortex, we characterized the encoding of melodic pitch, pitch-change, and expectation, and determined the extent to which these properties were encoded within separate or overlapping neural populations. The same participants also listened to natural speech, and we determined whether melodic feature encoding was specific to music or shared across domains.

## Results

To examine the neural encoding of melody, we created a naturalistic stimulus set consisting of 208 short musical phrases of varied instrumentation (Audio S1). Stimuli were designed to vary along three fundamental pitch-related dimensions (Fig. 1a): the absolute *pitch* (based on the fundamental frequency; F0) of each note, the *pitch-change* between adjacent notes, and the *expectation* of each note conditioned on prior notes in the phrase. Expectation was calculated using a pre-trained recurrent neural network (MelodyRNN; (*34*)) to estimate the surprisal of notes (negative log_2_ likelihood). Although these three measures are partially correlated, they contained sufficient independent variation for us to probe the extent to which they were independently encoded in the human auditory cortex (Fig. S1, Pearson’s correlation: pitch vs. pitch-change: *r* = 0.14; pitch vs. surprisal: *r* = -0.064; pitch-change vs. surprisal: *r* = 0.025).

Eight participants listened to musical stimuli while we recorded ECoG activity from high-density arrays placed over the lateral surface of the cortex. To identify music-responsive cortical sites, we extracted the average high frequency activity (HFA; 70 to 150 Hz) at each electrode during presentation of musical phrases (Fig. 1e). We observed activity primarily in bilateral STG, spanning posterior to mid-anterior subregions. Across all participants, we identified 224 electrodes with significant music-evoked responses relative to a silent baseline period (p < 0.01, Wilcoxon signed-rank tests, Bonferroni corrected).

### Distinct neural populations encode pitch, pitch-change, and expectation in melody

Next, we examined the extent to which music-responsive neural populations encoded relevant melodic information. At three example electrodes, we aligned cortical activity to the onset of each note (Fig. 1c) and compared evoked responses to notes that had high vs. low F0, were ascending vs. descending in pitch-change relative to the previous note, or were relatively expected vs. unexpected (median-split, independent two-sample t-tests, p < 0.001, Bonferroni corrected). At one electrode (Fig. 1c: left), responses differentiated notes in distinct pitch ranges, but did not differentiate notes with contrasting pitch-change or expectation values. At a different electrode (Fig. 1c: middle), responses differentiated descending from ascending pitch-changes, but not contrasts in either pitch or expectation. Finally, at a third electrode (Fig. 1c: right), responses differentiated how expected a note was but not its pitch or pitch-change. These response patterns demonstrate the sensitivity of local neural populations to distinct dimensions of melody (see Fig. S2 for all electrodes).

To quantify the encoding of these melodic dimensions at each electrode, we used temporal receptive field (TRF) modeling, which predicts continuous neural activity from a set of stimulus features (*35*). In addition to the three melodic features, we included the stimulus spectrogram and important temporal landmarks indicating phrase and note onsets in TRF models to statistically control for the cortical processing of features unrelated to – but potentially correlated with – pitch, pitch-change, and expectation (see Fig. S3 for full TRF predictor matrix).

Across electrodes, melodic and acoustic features explained a substantial portion of the variance in neural activity (max R^2^ = 0.35, mean R^2^ = 0.12), with the highest R^2^ values in an example participant located at cortical sites in the mid-to-anterior STG (fig. 2a). Performance was particularly high relative to the upper-limit defined by each electrode’s noise ceiling (*36, 37*), with models on-average predicting 70% of the explainable variance.

**Figure 2.**
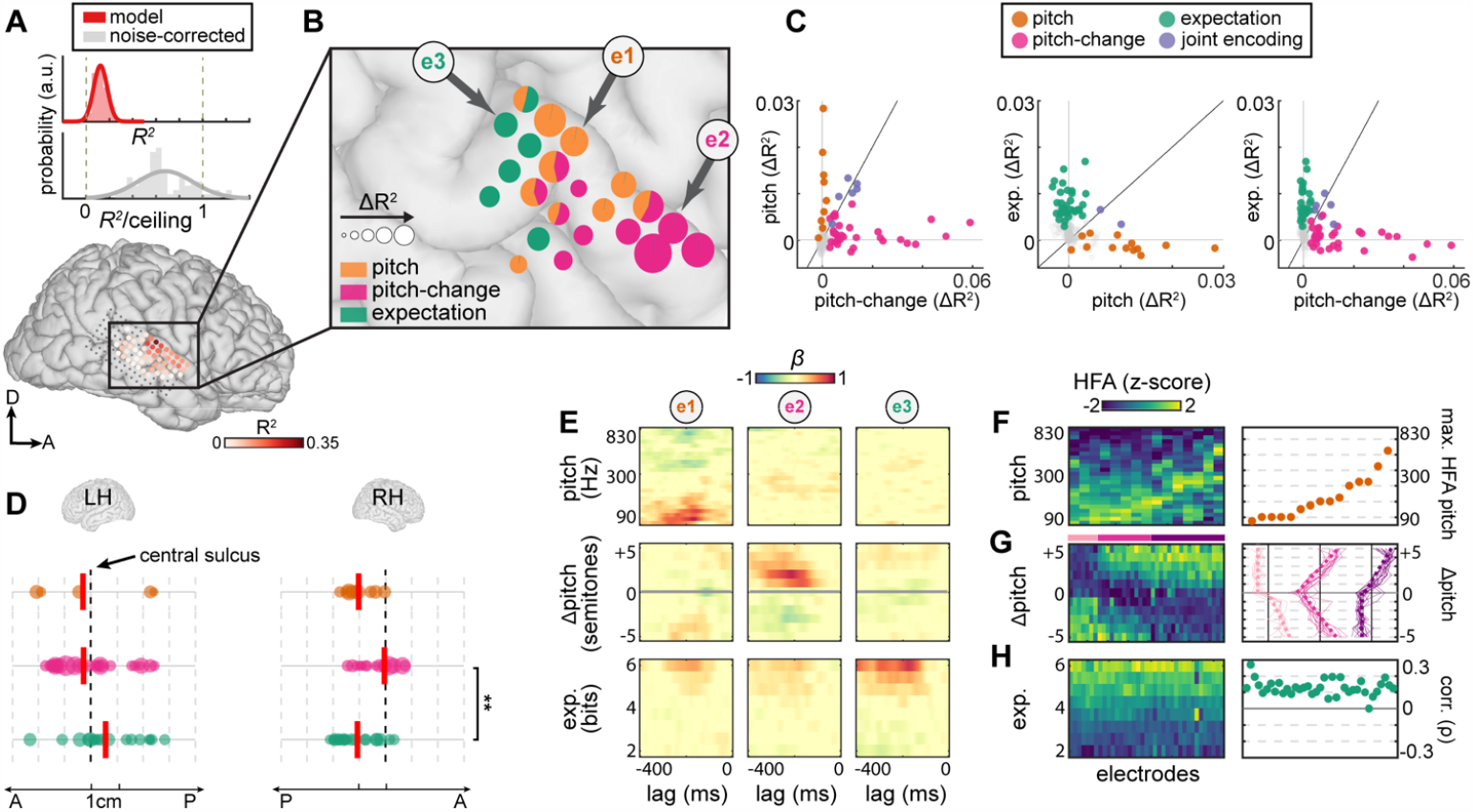
Separate neural populations encode melodic pitch, pitch-change, and expectation. **(A)** Top: Distribution of TRF model and noise-corrected *R*^*2*^-values across electrodes. Noise-corrected values are obtained by dividing model *R*^*2*^ by noise-ceiling estimates. Bottom: Electrode *R*^*2*^-values (indicated by darkness of markers) and neuroanatomical locations for an example participant. **(B)** Unique variance (Δ*R*^*2*^) explained by pitch, pitch-change, and expectation. Size of pie charts indicate the magnitude of Δ*R*^*2*^. **(C)** Scatter plots comparing Δ*R*^*2*^ explained by two given features. Electrodes are represented by markers and pooled across all participants, with colors indicating features with significant Δ*R*^*2*^ (permutation tests, p<0.01). **(D)** Electrode locations along the posterior-anterior axis of each hemisphere, normalized across participants to a common anatomical landmark. Marker size indicates normalized Δ*R*^*2*^. Vertical red lines indicate the weighted-mean location of feature encoding. Asterisks indicate significance level: ** p<0.01. **(E)** TRF weights for three example electrodes whose locations are shown in (B), showing tuning to low pitch (e1), ascending pitch-change (e2), and unexpected notes (e3). **(F)** Tuning to pitch. Matrix columns correspond to electrodes with significant ΔR^2^ explained by pitch and are sorted by peak F0. Right: Orange markers indicate F0s of peak response tuning. **(G)** Tuning to pitch-change. Electrodes are grouped by clusters (k-means, k=3) with colored bars above matrix indicating cluster membership. Right: Tuning profiles for electrodes within each cluster. Thin solid lines indicate individual electrode tuning. Thick dashed lines indicate linear functions fit separately on ascending and descending ranges of pitch-change. **(H)** Tuning to expectation. Right: For each electrode, green markers indicate rank-order correlations between HFA and expectation across all notes.

To determine the extent to which specific melodic features were encoded in single electrode activity, we computed the unique variance (ΔR^2^) explained by pitch, pitch-change, and expectation within TRF models (see methods). Within the STG of individual participants, we found electrodes that significantly encoded all three features (p<0.01, permutation tests). Crucially, encoding at single electrodes tended to be dominated by a singular melodic feature, with little-to-no encoding of the other two features (Fig. 2b, Fig. S3b).

Across all participants, the three melodic features were encoded within primarily nonoverlapping subsets of electrodes, with 80% of electrodes significantly tuned to a singular feature. Directly comparing the encoding of features across electrodes (Fig. 2c), we found that the ΔR^2^ attributed to any one feature was highly orthogonal to the ΔR^2^ attributed to the other two features (linear mixed-effects, random effects grouped by participant, all *β* not significantly different from zero, t-stat < 2.65, p > 0.05). Thus, in general, melodic features were not jointly encoded within the same neural populations. Rather, pitch, pitch-change, and expectation were selectively encoded within three distinct subpopulations in the STG.

We next asked whether these three distinct subpopulations were anatomically organized. We found that melodic features were represented within highly overlapping regions along the posterior-to-anterior axis of the STG (Fig. 2d; see Fig. 2b and Fig S3b for organization within individual participants). In the right hemisphere, the distribution of pitch-change encoding electrodes was significantly more anterior to the distribution of expectation encoding electrodes (linear mixed effects, randomized block design, Bonferroni-corrected: t-stat = 3.50, p = 0.0056). Despite this coarse spatial difference, the representation of melodic features did not strongly segregate into anatomically distinct auditory subregions.

Beyond determining *whether* auditory cortical populations encode important dimensions of melody, we were interested in *how* this information is coded. To answer this, we first inspected model weights for three example electrodes (Fig. 2e) that had significant ΔR^2^ explained by pitch (e1), pitch-change (e2), and expectation (e3). Consistent with their corresponding ΔR^2^ values (Fig. 2c), weights were concentrated over a single feature at each electrode, and revealed clear patterns of tuning to low pitch at *e1*, ascending pitch-changes at *e2*, and more unexpected notes at *e3*.

To further characterize the specific format in which pitch was coded, across all electrodes with significant ΔR^2^ explained by pitch, we characterized patterns of response-tuning by visualizing the modulation in activity across the F0 range spanned by the stimulus (Fig. 2f). This revealed a diversity of broad tuning profiles, with maximal responses tiling the F0-range from low (90Hz) to mid-high (300-700 Hz) values.

While we used F0 as a proxy for pitch, the perception of pitch is influenced by several other acoustic properties that are correlated with F0 (*38, 39*). This includes the spectral profile of notes (linear correlation, spectral centroid vs. F0, *r* = 0.75) and spectral modulations (modulation centroid vs. F0, *r* = -0.4). We therefore asked whether such alternate spectral properties could explain current pitch encoding above and beyond F0. First, we compared F0 tuning across different subsets of the musical stimulus with highly distinct spectral profiles (Fig. S4). Tuning curves were strongly correlated across spectrally distinct stimuli (t>3.4, p < 0.001). Next, we used a variance partitioning approach to determine whether spectral modulations could explain neural responses better than F0. Model R^2^ values were significantly higher when we maintained F0 as a proxy for pitch, rather than spectral modulations (one-sample t-test, *t* = 4.6; p = 3.9×10^−4^). Additionally, F0 continued to explain ΔR^2^ even when controlling for spectral modulations (Fig. S5). These results indicate that, while pitch perception is influenced by multiple acoustic attributes, the observed STG encoding of pitch is driven by neither the spectral profile of notes nor their spectral modulations, suggesting mechanisms more directly related to the extraction of F0.

Next, to further characterize encoding of pitch-change, we visualized response modulation across the range of F0-changes in the stimulus (fig. 2g, fig. S8a). Prior behavioral research has suggested separability in the representation of the precise magnitude of pitch-changes (interval) versus their general direction (contour) (*10, 12*). However, we found local tuning patterns that represented both contour and interval information. Most electrodes (63%) had rectified-linear tuning profiles, with responses selective for a given direction (ascending or descending) and positively modulated by the magnitude of change within the preferred direction (Fig. 2g; light pink and purple clusters, k-means clustering). Additionally, we found electrodes that responded proportionally to interval-magnitude, regardless of direction (Fig. 2g, magenta cluster). Thus, STG populations tuned to pitch-change represented both melodic contour and interval.

Importantly, psychophysical evidence suggests that the perception of pitch-change in humans is mediated by multiple mechanisms, with listeners relying on comparisons between the F0 as well as the individual harmonic components between adjacent notes (*38, 40*). We next examined the extent to which the observed neural encoding reflected these two possibilities. First, we characterized response tuning separately within two distinct subsets of the data, grouped by whether higher-order harmonic components provided an unambiguous versus ambiguous cue to the direction of change. In both subsets, tuning remained the same (Fig. S7). To further dissociate harmonic cues from those based on the F0, we created two sets of artificial melodic stimuli. One set was comprised of harmonic complex tones, while another comprised tones in which higher-order components were jittered, rendering them inharmonic and thus lacking a reliable F0. We identified an electrode at which activity was modulated by pitch-change in both original melodies and harmonic stimuli. Crucially, activity at this electrode was not modulated by pitch-change for inharmonic stimuli (Fig. S6). These findings suggest the presence of auditory populations that encode pitch-change by primarily tracking F0-changes. Further work is needed to identify the neural basis of pitch-change detection based on tracking of harmonic components, and to understand how these multiple representations contribute to our ultimate perception of pitch-change.

Finally, we examined the encoding of expectation, which reflects the degree to which successive notes in the melody conformed to or departed from sequential patterns and learnt structural rules of Western tonal music (Fig. S9). Across electrodes with significant ΔR^2^ explained by expectation, we consistently found a monotonic relationship, whereby more unexpected notes evoked stronger responses (Fig. 2h).

These results demonstrate that, in higher-order auditory cortex, perception of melody recruits multiple anatomically and functionally independent subpopulations, each selectively tuned to information along a different pitch-related dimension, spanning basic spectral to time-integrated and statistical structure.

### Music-selective activity reflects encoding of melodic expectation

Beyond identifying how relevant information is encoded, a major goal of auditory neuroscience is to determine the nature and extent of specialization in the human brain for music, compared with other acoustically complex and behaviorally relevant sounds like speech. Recent work has found subregions in the STG that are selectively activated by music over other sounds (*31–33*), suggesting domain specialization. To date, however, the dimensions of music to which these subregions are selectively tuned remains unclear.

To address this question, we presented the same eight participants with naturally spoken English sentences (Audio S2). We hypothesized that music-selectivity reflects tuning to information that exclusively exists within music. Specifically, while pitch and pitch-change are acoustic properties that describe general pitch-based information that exists across different domains, melodic expectation describes the unique sequence structure of music. We therefore predicted that the degree to which populations selectively respond to music (over speech) directly reflects the extent to which they encode expectation.

To identify music selective electrodes, we first compared the relative magnitude of music and speech responses (fig. 3a). From this, we derived a selectivity index (SI) ranging from -1 (speech selective) to +1 (music selective) quantifying the degree to which a given electrode preferentially responded to a given domain. While most electrodes responded to both music and speech, we nevertheless identified a substantial number of electrodes that were consistently and strongly selective for musical phrases over spoken utterances (fig. 3b).

**Figure 3.**
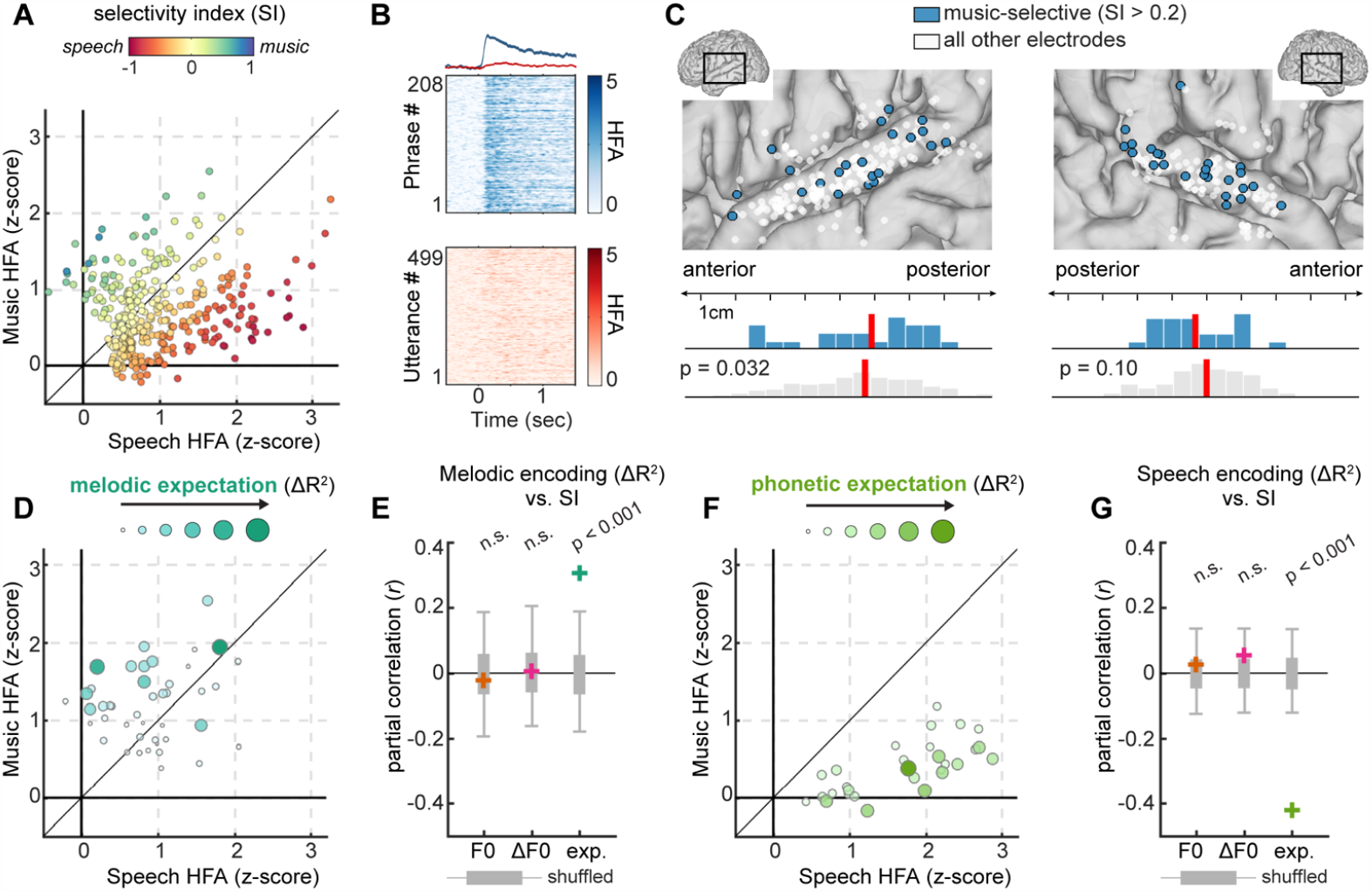
Music selective activity reflects the encoding of melodic expectation. **(A)** Average music versus speech responses for all electrodes. Marker colors indicate selectivity index (SI). **(B)** Single trial rasters for an example electrode that demonstrates selective responses to music (blue) over speech (red). **(C)** Anatomical location of music-selective electrodes (SI > 0.2; blue markers) indicating broad distribution throughout STG. Histograms indicate distribution of music-selective electrodes relative to all other electrodes along the posterior-anterior axis. Vertical red lines indicate median location of distributions. **(D)** Average music versus speech responses for electrodes that encode melodic expectation (axes are identical to panel A). Marker size and color indicate ΔR^2^ explained by expectation in TRF models. **(E)** Colored crosses indicate the partial correlation between SI and encoding of pitch (orange), pitch-change (magenta) and expectation (dark green) across all electrodes with SI ≥ 0. Gray error bars indicate 95-percentiles of permutation tests. **(F)** Average music versus speech responses for electrodes that encode phoneme-based expectation in speech. Marker size and color indicate ΔR^2^ explained by phonetic expectation in TRF models of speech-evoked activity. **(G)** Partial correlation between SI and speech encoding across all electrodes with SI ≤ 0.

Anatomically, music selective electrodes (defined as electrodes with SI > 0.2, see Methods) were widely distributed and intermixed with other sound-responsive populations in the STG (fig. 3c). The music selectivity of electrodes was independent of location in the right hemisphere (linear mixed-effects, randomized block design, Bonferroni corrected: t = 1.97, p = 0.13) and weakly biased towards posterior regions in the left hemisphere (t = 2.45, p = 0.032). Further, within-subject comparisons revealed only one participant in which the spatial distribution of music selective electrodes differed from that of nonselective electrodes (ranksum tests, p = 0.014 in one participant, all other p > 0.05). Thus, we find little evidence for the clustering of music selectivity into a dedicated auditory subregion.

While the existence of music selective populations is consistent with prior studies (*31*), we next sought to evaluate our key hypothesis – that music selectivity specifically reflects the encoding of melodic expectation. We visualized the strength of expectation encoding (ΔR^2^) as a function of selectivity (Fig. 3d). Consistent with our hypothesis, electrodes that encoded expectation were almost exclusively more responsive to music than speech. Furthermore, across all music-selective electrodes, the degree of expectation-encoding predicted the magnitude of music selectivity (fig. 3e; partial correlation, *r* = 0.31, p < 0.001, permutation test). In contrast, there was no systematic relationship between selectivity and the encoding of pitch or pitch-change (pitch: *r* = -0.02, p = 0.6, contour: *r* = 0.006, p =0.47. fig. S10).

As linguistic sequences can also be characterized by their statistical structure, and prior studies have shown robust encoding of phoneme-based expectation in the STG (*41, 42*), we next sought to test the analogous hypothesis for populations that were speech selective. Using the same TRF modeling approach as before, we quantified the encoding of a set of acoustically and perceptually relevant speech features in sentence-evoked activity (see methods for details). Mirroring music, we found that phoneme-based expectation was encoded by speech selective electrodes (Fig. 3f), with the degree of encoding predicting the magnitude of speech selectivity (Fig. 3g; expectation: *r* = -0.42, p < 0. 001, pitch: *r* = -0.026, p =0.65, contour: *r* = 0.054, p =0.77). Thus, rather than a domain-general mechanism for representing auditory sequence statistics, the relevant statistics of music and speech are encoded within two independent substrates of the STG.

So far, results suggest that selectivity for music reflects the encoding of its higher-order sequence structure, rather than its lower-order acoustic properties. However, music has an acoustic structure that is inherently different from that of speech, particularly in terms of the spectral and temporal modulation patterns to which STG populations are sensitive (*43–46*). We therefore sought to explicitly test whether low-level acoustic differences – rather than expectation – explain music selectivity. To do so, we created a third stimulus that was acoustically identical to speech except that it contained the pitch structure of melody (Fig. 4a-c; Audio S2 and Audio S3). Specifically, for every speech utterance, we created a “melodic-speech” counterpart by warping the continuous pitch within each syllable onto the nearest discrete Western musical-scale tone. Importantly, this manipulation left all other acoustic features, such as vowel formants and amplitude envelopes, unchanged (compare Fig. 4a-b). As a result, the spectrograms of speech and melodic-speech signals were virtually identical, with correlations close to 1 along both the spectral (*r* = 0.998) and temporal (*r* = 0.995) axis. For a subset of original sentences (118/499), their melodic-speech counterparts formed coherent Western diatonic melodies, which we determined using a musical key-finding algorithm (*47, 48*).

**Figure 4.**
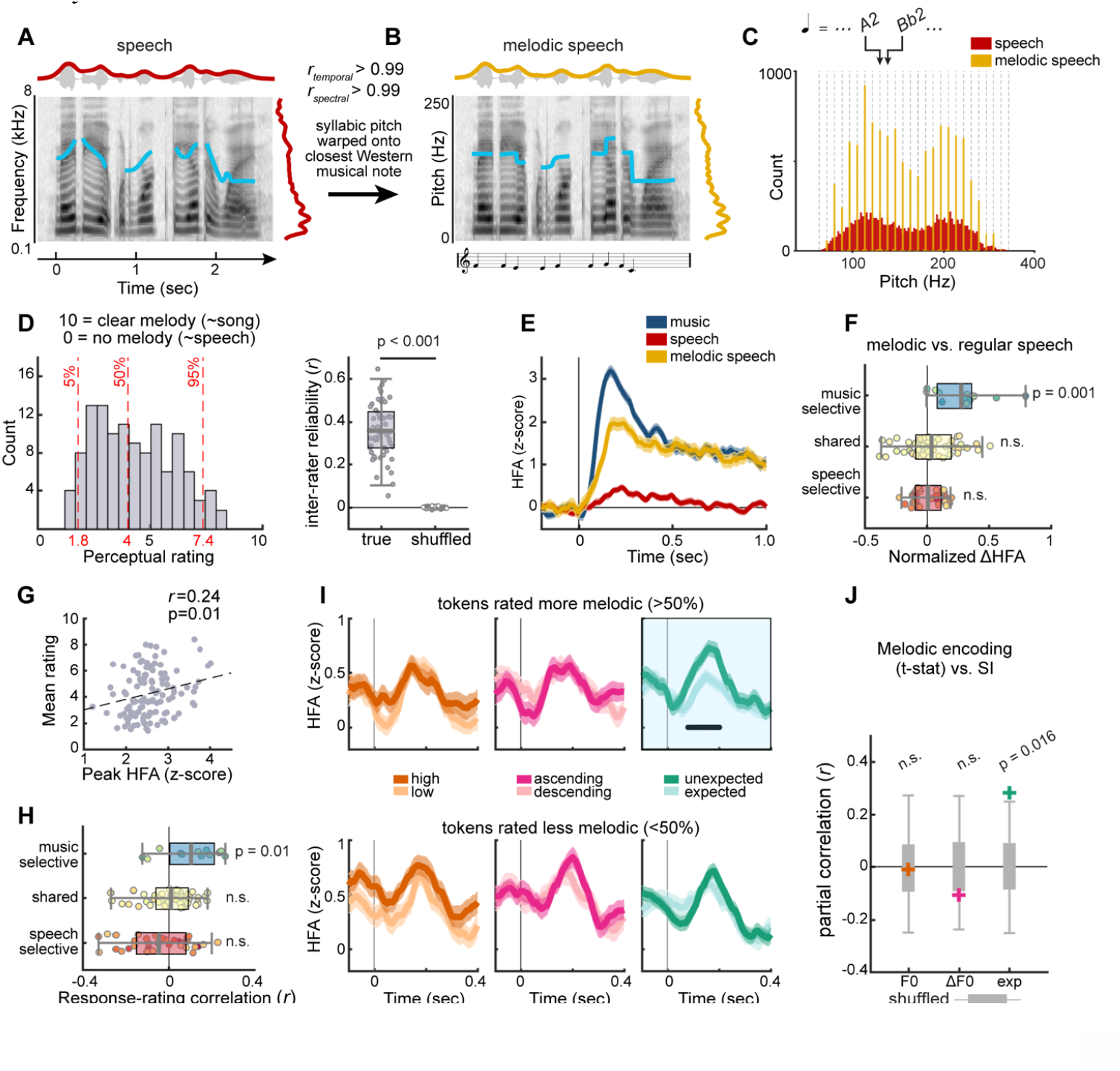
Music-selectivity is independent of low-level spectrotemporal properties. **(A)** Spectrogram of an example speech token. Overlaid blue lines indicate pitch contour. Dark red lines above and to the right indicate temporal and spectral envelopes respectively. **(B)** Melodic-speech spectrogram for the same token as in (A), illustrating a near identical spectrotemporal structure to speech. Musical notation underneath indicates the discrete musical pitch of each syllable. **(C)** Pitch distributions show that melodic-speech discretizes syllabic pitch to the nearest Western musical note. **(D)** Left: Distribution of mean ratings indicating the extent to which independent listeners (N=11) heard each melodic-speech token as melody. Red dashed lines indicate 5^th^, 50^th^, and 95^th^ percentiles of distribution. Right: Inter-rater reliability. **(E)** Mean evoked responses to music, speech, and melodic-speech at a music-selective electrode. **(F)** Response difference indicating the extent to which melodic-speech elicited larger responses than speech across electrodes split into speech-selective (SI < -0.2), shared (−0.2 < SI < 0.2) and music-selective (SI > 0.2) bins. Marker colors indicate electrodes’ SI. **(G)** Scatter plot showing correlation between the peak response to melodic-speech (x-axis) and perceptual ratings (y-axis) across all tokens for the same electrode as panel E. **(H)** Response-rating correlations across all electrodes split into the same selectivity-dependent bins as in panel (F). **(I)** Tuning to melodic features within melodic-speech for the same music-selective electrode as (E). Separate traces indicate tuning for tokens perceived more (top row) and less (bottom row) like melody. Responses are grouped based on the median-split along each feature dimension. **(J)** Colored crosses indicate partial correlations between SI and the modulation of melodic-speech responses by melodic features. Gray bars indicate the 95-percentile of permutation tests.

To evaluate the perception of melodic-speech, an independent cohort of listeners (N=11) rated the extent to which they heard a melody within each token on a scale ranging from 0 (sounds like regular speech) to 10 (sounds like song). We found that all tokens were perceived to contain melody, albeit with wide variation in the degree to which this was the case (Fig. 4d). Furthermore, ratings were highly consistent across listeners (average inter-rater reliability: *r* = 0.35, p =2.3 × 10^−27^, permutation test), suggesting that they served as a reliable proxy for the degree to which ECoG participants experienced melodic-speech as melody.

A subset of ECoG participants (N=2) who previously heard music and speech stimuli were also presented with melodic-speech. At music-selective electrodes, we first asked whether melodic-speech elicited similar responses to music. If so, this would specifically implicate information imparted by the morphing of original speech into discrete musical tones – and not other spectrotemporal features – as the basis of music selectivity. For an example music-selective electrode, melodic speech elicited robust responses that were qualitatively similar to those evoked by music, and stronger than those evoked by speech (Fig. 4e). Such an enhanced response to melodic versus regular speech was primarily observed at music selective electrodes (Fig. 4f; music-selective: p = 0.0011; shared: p=0.13; speech-selective: p=0.39, one-sample t-tests) and was positively correlated with electrodes’ SI (*r* = 0.66; p = 1.8×10^−6^). Thus, inserting the pitch structure of melody into a stimulus that otherwise had the spectrotemporal characteristics of speech was sufficient to elicit music-like responses at music-selective electrodes.

We next asked whether enhanced responses to melodic (versus regular) speech specifically arose from the encoding of melodic expectation. Because listeners were unlikely to develop melodic expectations for stimuli not strongly perceived as melody, we first asked whether melodic-speech responses scaled with perceptual ratings. Indeed, we observed a positive correlation between ratings and responses both at an example electrode (Fig. 4g; linear correlation: *r* = 0.24; p = 0.01) and across music-selective electrodes (Fig. 4h; one sample t-tests: music-selective p = 0.011; speech-selective p = 0.97; shared p = 0.29). This implies that music selective responses are driven by information that exists only to the extent that stimuli are perceived as melody.

Finally, to explicitly test whether responses to melodic speech were modulated by features of melody, we extracted the pitch, pitch-change, and melodic expectation of each token (using melodyRNN to extract expectation). We divided each feature distribution into two equal bins (median-split) and plotted the corresponding neural responses in each bin. At an example music-selective electrode, responses to melodic speech were significantly modulated by expectation (independent two-sample t-test: p < 0.05). However, we only observed this modulatory effect for tokens that induced a relatively strong percept of melody (Fig. 4i; top vs. bottom row). In contrast, responses were not modulated by pitch or pitch-change (independent two-sample t-test, p>0.05). Mirroring natural music, the degree to which expectation modulated melodic-speech activity predicted electrode SI, while modulation due to pitch or pitch-change did not systematically vary with selectivity (Fig. 4j; partial correlation: pitch, *r* = -0.011, p = 0.55; pitch-change, *r* = -0.11, p = 0.79; expectation, *r* = 0.28, p = 0.016, permutation tests). Taken together, the above results indicate that, for both natural music and melodic-speech alike, domain-selective activity specifically reflects the encoding of melodic expectation.

### Encoding of pitch and pitch-change is shared across music and speech

Having established that the encoding of melodic expectation is functionally specialized for music, we next probed whether lower-level dimensions of melody – pitch and pitch-change – are represented by domain-general neural codes. First, we characterized speech along dimensions that are acoustically equivalent to melodic pitch and pitch-change. To do so, we extracted the pitch contour of each sentence and computed the suprasegmental changes between the median pitch of adjacent syllables (Fig. 5a-b). Although not identical, distributions of pitch and pitch-change were highly overlapping across the two domains (Fig. 5c).

**Figure 5.**
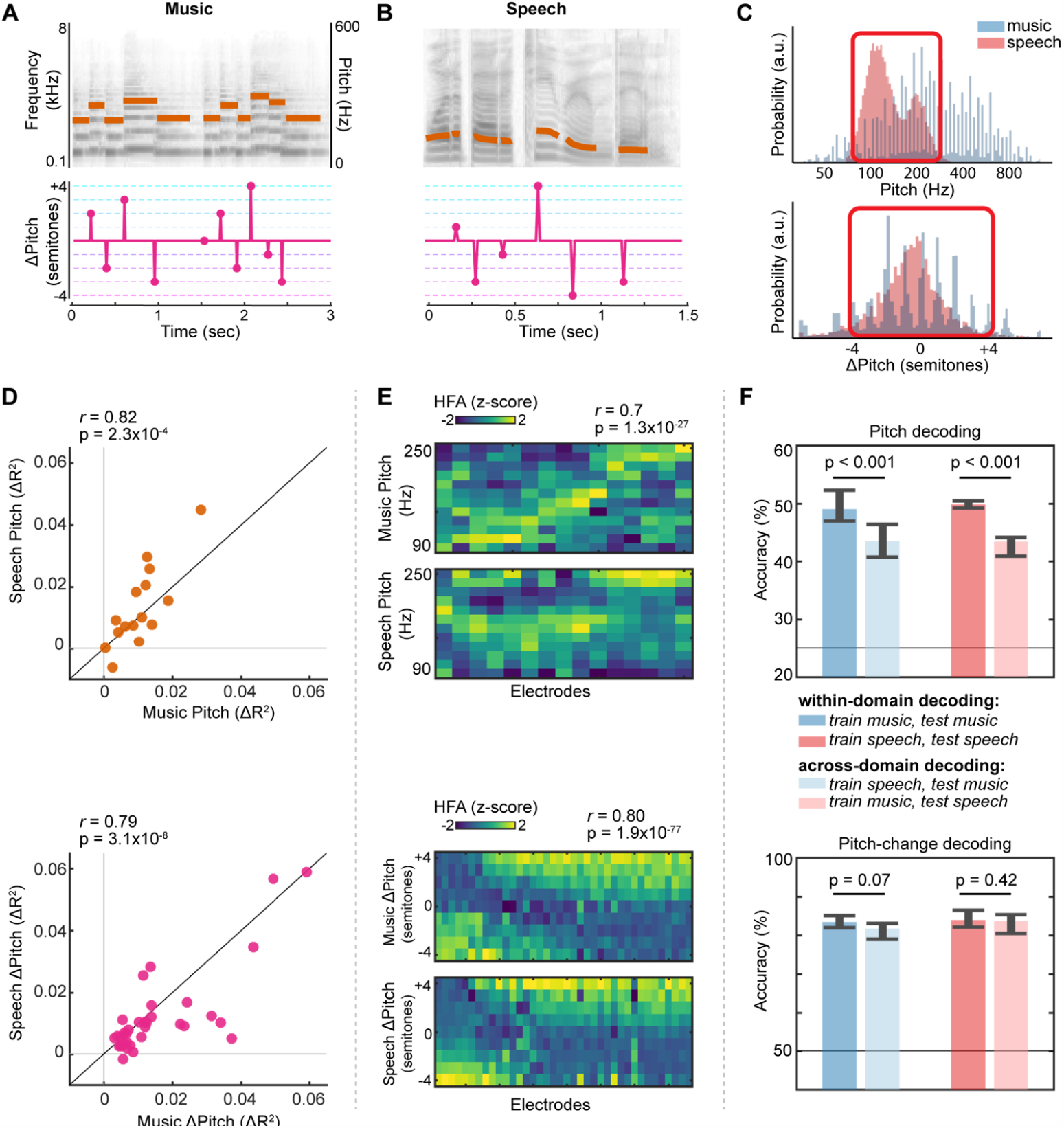
Shared representations of pitch and pitch-change across music and speech. **(A-B)** Pitch and pitch-change representations for example music and speech tokens. Orange lines indicate pitch contours overlaid on stimulus spectrograms. Pitch-change in speech is based on differences in median syllabic pitch, specified at syllable onsets. **(C)** Partially overlapping pitch (top) and pitch-change (bottom) distributions in music and speech. Red boxes indicate overlapping regions of feature space. **(D)** Δ*R*^*2*^ explained by pitch (top) and pitch-change (bottom) in TRF models of music (x-axis) versus speech (y-axis). **(E)** Tuning curves for pitch (top) and pitch-change (bottom) within music and speech. Tuning is characterized for every pitch or pitch-change encoding electrode (x-axis) across the overlapping range of feature-space found in music and speech (y-axis). For pitch, columns are ordered by the F0 corresponding to peak-HFA in music. For pitch-change, columns are ordered by increasing HFA difference between ascending and descending changes. **(F)** Linear classification accuracy when decoding pitch (top) or the direction pitch-change (bottom) from corresponding neural activity across electrodes. Classifiers are trained and tested on neural activity either within (darker colored bars) or across domains (lighter colored bars). Errors indicate 95% distribution of bootstrap tests. Horizontal black lines indicate chance accuracy.

Next, for electrodes that significantly encoded pitch or pitch-change in melody, we examined the extent to which they encoded information along the same dimension of speech. As with music, we computed the unique variance (Δ*R*^*2*^) explained by each feature within TRF models of speech-evoked activity. We then directly compared Δ*R*^*2*^ values across the two domains (Fig. 5d). We found strong correlations in the extent to which electrodes encoded either pitch (*r* = 0.82, p = 2.3×10^−4^) or pitch-change (*r* = 0.79, p = 3.1×10^−8^), indicating that neural populations strongly tuned to a given dimension of melody were generally tuned to the same dimension of speech.

While the above findings suggest that the same populations encode pitch and pitch-change across domains, we next probed the extent to which these populations represented information using a domain-general neural code. We computed tuning curves for the overlapping range of pitch and pitch-change across music and speech. This overlapping range spanned from 80 to 265 Hz for pitch, and -4 to +4 semitones for pitch-change (Fig 5c). Both within and across electrodes, tuning profiles were highly correlated across domains for pitch (Fig. 5e, top, *r* = 0.70, p = 1.6×10^−27^) and pitch-change (Fig. 5e, bottom, *r* = 0.82, p = 1.6×10^−74^), indicating a highly conserved neural code across music and speech.

To further quantify the extent of domain generalization, we first trained linear classifiers to decode pitch (Fig. 5f, top) or the direction of pitch-change (Fig. 5f, bottom) from electrode activity pooled across participants. We trained separate classifiers to decode information in music and speech activity. We then tested these classifiers using activity from either the same domain (darker bars) or the opposite domain to the one in which they were trained (lighter bars). As expected, within-domain decoding accuracy was well above chance for both features (bootstrap tests, all p < 0.001). Crucially, when decoders were tested across domains, accuracy remained well above chance for both pitch and pitch-change (bootstrap tests, all p < 0.001). In particular, for pitch-change, within and across domain decoding did not significantly differ (bootstrap difference tests: music generalization, p = 0.07; speech generalization, p = 0.42). Thus, in contrast to the encoding of melodic expectation, lower-level properties of melody are represented by general-purpose auditory populations that largely share a neural code with speech.

## Discussion

Despite decades of neuroimaging studies implicating a role for the STG in perception of music (*49–52*), until now the specific information represented in this region has remained unclear. Using high-density direct recordings from human auditory cortex, we demonstrated evidence for the extraction of multiple perceptually important pitch-based features of melody. We showed that the encoding of one such feature – melodic expectation – explains activity in populations that selectively respond to music. In contrast, shorter timescale acoustic features are encoded by general-purpose auditory populations.

Despite all relating to pitch, different features of melody were selectively encoded within separate neural populations, consistent with a model in which distinct types of information are processed in dissociable pathways (*19–21*). This may arise via parallel projections from earlier cortical or subcortical regions (*53*). Future work should explicitly identify the inputs to each subpopulation to determine the network architecture underlying the observed spatial code for melody.

Current findings clarify the nature and extent of specialization for music in the human brain. Previous research has shown the existence of music-specific cortical activity (*31–33*). While evidence suggests that this activity is not explained by low-level sensory properties (*46*), it has remained unclear which properties of music do in-fact drive responses in these populations. Here, we found that music-selectivity is systematically driven by the encoding of melodic expectation in a format consistent with predictive coding theory, whereby more unexpected notes evoke larger responses (*18, 54–56*). Our results demonstrate functional specialization in the human brain for encoding the sequential structure of behaviorally relevant classes of sound. Future work is needed to determine the degree to which this encoding is “bottom up”, reflecting regularities within recent stimulus history, versus “top-down”, potentially requiring feedback from higher-order cortical regions (*57*).

Finally, current results reveal the extent to which auditory representations are shared across music and speech. Neurophysiological work in non-human animals has characterized the encoding of pitch-based features using artificial stimuli that are domain-agnostic (*58, 59*). More recently, human electrophysiology has examined perceptually relevant pitch-based features of speech (*19, 60*). Despite these advances, the extent to which music and speech perception leverage a shared neural architecture for representing overlapping acoustic information has been unclear. Evidence from behavioral and lesion studies and has been conflicting, with some studies suggesting shared neural mechanisms (*27, 61, 62*) while others suggest anatomically and functionally separable systems (*63*). Leveraging the spatiotemporal resolution of ECoG, we provide direct evidence that, even within higher-order auditory regions, representations of musical pitch and pitch-change are largely domain-general. These representations were broadly localized to anterior regions of the STG, consistent with prior reports of pitch-sensitive cortical areas (*64*). For the purpose of comparing speech with music, we coded pitch within speech as a discrete value based on the median F0 within each syllable. However, future work must examine the domain-generality of encoding mechanisms that support the perception of continuous pitch variation, such as that occurring within-syllables or across more diverse musical genres.

The extraction of multiple pitch-based dimensions is central to our perception of melody. Our findings underscore the multidimensional nature of this encoding in the human brain – that is, melody is not processed by a single pitch-sensitive cortical area. Rather, distinct populations encode progressively higher-order pitch representations across the auditory cortex, making use of both general-purpose and music-specific mechanisms to do so.

## Materials and Methods

### Participants

Eight patients undergoing treatment for intractable epilepsy participated in the study (see table S1 for demographic and clinical information). One additional patient participated in a control experiment involving presentation of harmonic or inharmonic complex tones (see below) and was also presented with a partial dataset of music (3/6 blocks). Participants were implanted with 4mm-spaced subdural electrode grids unilaterally over peri-Sylvian regions for clinical monitoring of seizure-related activity. Grid placement was determined solely by clinical considerations. One patient reported a history of tinnitus while all others reported having normal hearing. All patients were non-musicians with the extent of prior formal musical training ranging from 0 – 8 years (table 1). Power analyses were conducted using g-power (https://www.psychologie.hhu.de/) based on the expected mean and standard deviation in melodic feature encoding. Required sample sizes suggested that the recruitment of 8 participants provided sufficient statistical power for analyses.

### Software

Analyses were carried out using custom-written scripts in MATLAB 2016b (MathWorks, www.mathworks.com) and Python. Cortical surface reconstruction was performed using Freesurfer and electrodes were localized using a Python package (img-pipe). Melodic expectations were extracted using publicly available Python toolboxes (https://github.com/magenta/, (*34*) and all other melodic features were extracted in MATLAB using the music information retrieval toolbox (*65*) and custom-written scripts. Manipulation of pitch for creating melodic speech stimuli was performed in Praat (*66*). Syllable onsets and offsets in speech were detected using a forced aligner (https://babel.ling.upenn.edu/phonetics).

### Stimuli and procedure

All participants passively listened to natural music and speech stimuli while we recorded ECoG activity. Two of these participants additionally passively listened to a control stimulus which we refer to as melodic speech. All stimuli were delivered through free-field speakers from a Windows Laptop at a sample rate of 44.1kHz (melody) or 16kHz (speech and melspeech).

#### Music

A stimulus set comprising 214 distinct monophonic musical phrases (total duration = 23.4 minutes, mean duration = 6.57 seconds) was compiled by sampling directly from natural solo instrumental recordings. We excluded six purely percussive phrases from the current analysis. The remaining phrases cumulatively contained 4578 discrete notes, with a mean density of 3.4 notes per second. Phrases collectively featured 18 different Western instruments and a variety of genres broadly categorized as classical, jazz, or folk (see table 2 for example source material). Phrases were presented in pseudorandom order and separated by silent inter-trial intervals ranging from 0.7 – 1.5 seconds. Participants heard each phrase once, with data collected over 5 separate listening blocks lasting approximately 4 minutes each. An additional block contained 10 repetitions of 10 phrases. Musical phrases were chosen to avoid well-known melodies, and participants reported being unfamiliar with most phrases.

#### Speech

Speech stimuli comprised a selection of 499 English sentences from the TIMIT corpus (*67*), spoken by a variety of male and female speakers with regional North American accents. Speech stimuli were presented to participants in a similar fashion to music – in pseudorandom order and over 4 separate listening blocks with an additional block containing 10 repetitions of 10 sentences. Sentences had a mean duration of 2.05 seconds (SD = 0.4) and were presented in pseudorandom order.

#### Melodic-speech

Melodic-speech control stimuli were created by altering the pitch of TIMIT tokens while leaving all other spectrotemporal features of speech intact. To create melodic speech, we first calculated the median pitch of each syllable and identified the closest Western-musical pitch. We then warped each syllable’s continuous pitch onto its nearest discrete musical value, thereby transforming each syllable into a discrete musical pitch event (fig. 4a-c). To verify that this process did not significantly change the spectrotemporal structure of speech, we compared the spectral and temporal profiles of melodic speech and speech. Specifically, to compare temporal structure, we averaged power across all frequency bands of the spectrogram to extract the temporal envelope for every token. We concatenated all tokens into two vectors for speech and melodic-speech, respectively, and computed their linear correlation. As expected since melodic speech is derived from speech, temporal envelopes were nearly identical (r = 0.995). To compare spectral structure, we concatenated power across frequency bands at every timepoint in the spectrograms into two vectors for speech and melodic-speech respectively. Spectral envelopes were strongly correlated (r = 0.998). This was expected since speech and melodic speech only differed in that the latter had a discretized within-syllable F0. To evaluate whether this modification to speech generated coherent melodies, we first applied an automatic musical key finding algorithm to each token (*47, 48*). This algorithm returns a series of 24 correlation coefficients indicating the extent to which the sequence of pitches aligns with the canonical pitch distributions of the 24 Western diatonic keys (12 major, 12 minor). We retained tokens with a maximum correlation coefficient greater than 0.6, resulting in 118 melodic speech tokens. For comparison, the mean output of the key finding algorithm applied to original musical stimuli was 0.77 (standard deviation = 0.11). We presented melodic-speech tokens grouped by their tonality, with each new tonality primed by a piano chord that outlined the root of the new key. The duration of the entire stimulus was 3 minutes, which a subset of 2/8 ECoG participants listened to twice.

#### Harmonic and Inharmonic Tones

For a subset of the original music stimulus (47 out of 214 phrases), we used the estimated F0 and note-durations of melodies to generate synthesized tone-sequences. This subset was pseudo-randomly chosen such that it contained pitch and pitch-change ranges representative of the entire musical stimulus. We synthesized two versions of each melody, one consisting of harmonic tones and the another consisting of inharmonic tones. Each tone consisted of 6 harmonic components including the F0 with the relative power of each component preserved from original melodies. We applied a tapered cosine window to each tone with 10ms onset and offset ramps. To create inharmonic tones, we followed the identical procedure to that used in (*38, 40*). Specifically, we jittered the frequency of each harmonic, excluding the fundamental, by a random amount chosen from the uniform distribution U(−0.5, 0.5). This value was multiplied by the F0 and added to the frequency of the respective harmonic. To reduce beating, jitter values were further constrained such that all frequency components were separated by at least 30Hz. For a given melody, the same profile of jitter values was applied to every note.

### Melodic feature extraction

Note onset times and their corresponding absolute pitch values were extracted using the Music Information Retrieval toolbox (*65*) and verified manually. Pitch-change was then calculated by subtracting the pitch value of the previous note from that of the current note, and expressed in semitones. To extract melodic expectations, we used a Recurrent Neural Network model (MelodyRNN, https://github.com/magenta/) that applies natural language modeling approaches to model melodic sequence structure. We used an “off-the-shelf” implementation in which the model was pre-trained on a large corpus of Western melodies with the following parameters: learning rate = 0.001, batch size = 128, number of layers = 2 × 512 nodes, attention length = 40, dropout rate = 0.5. Internal model weights were optimized during training to maximize the probability mass assigned to the note occurring at *t*_*n+1*_ given the pitch and duration of previous notes from *t*_*1:n*_.We used the “Attention” configuration of melodyRNN (*34, 68*), which enables the model to learn long-term dependencies characteristic of music that traditional Markov-based approaches fail to capture (*69*). For all phrases in the current stimulus set, melodyRNN was used to calculate the surprisal of each note, defined as the negative log probability of the note *e* that occurred at position *i*, given the MIDI pitch and duration (quantized to the nearest 16^th^ note) of preceding notes in the melody:

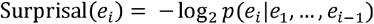

We also computed the uncertainty of the melody at each timestep, which is defined as the entropy of the probability distribution over an alphabet of *K* possible notes:

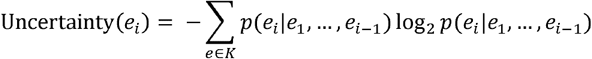

Surprisal and uncertainty are complementary measures in that the former indicates the extent to which an event deviated from preexisting expectations, while the latter conveys the specificity of those expectations in anticipation of the event. While these measures are highly correlated (r = 0.32, p < 0.001; Fig. S1) and produced similar patterns of neural tuning (Fig. S3C), we found that surprise modulated evoked responses to a greater degree than uncertainty (Fig. S2). While melodyRNN has been found to generate realistic melodies (*34, 68*), to sought to further validate its ability to accurately model listeners’ expectation. We tested its ability to correctly predict the next note in a set of Western melodies and found high accuracy (> 90%). We also computed expectation using a traditional Markov approach (*70*), and found correlations between the two different modeling approaches (surprise: *r* = 0.59, p < 0.001; uncertainty: *r* = 0.38, p < 0.001). We found that melodyRNN explained a greater amount of variance in neural activity than Markov-based estimates (t = 4.48; p = 1.01×10^−5^), which may reflect its ability to model long-distance dependencies as noted above.

### Neural recordings and preprocessing

Electrocorticographic (ECoG) activity was acquired at a sampling rate of 3051.8 Hz using either a 256 channel PZ2 amplifier or 512 channel PZ5 amplifier connected to an RZ2 digital acquisition system (Tucker-Davis Technologies, Alachua, FL, USA). We recorded the local field potential of each electrode and removed line-noise using notch-filters at 60 Hz, 120 H, and 180 Hz. Bad channels with variance indistinguishable from noise or continuous epileptiform activity were removed, and time segments on remaining channels that contained electrical or motor artifacts were marked and excluded. We used the log-analytic amplitude of the Hilbert transform to filter the signal into eight log-spaced bands in the high-gamma range from 70-150 Hz and took the first principal component across these bands to extract stimulus-related activity (*71, 72*). The resulting high-frequency activity (HFA) was downsampled to 100Hz. When fitting TRF models, we normalized activity by subtracting the mean and dividing by unit variance (i.e., the z-score) using activity across entire recording blocks. In general, for naturalistic stimuli like music, we have found that this normalization approach yields higher TRF performance and more stable weights than normalizing to a local baseline. However, for analyses involving the comparison of activity *across* stimulus domains (for instance, when deriving a domain selectivity index), signals were re-normalized relative to the mean and standard deviation of a 500ms silent period preceding each individual token. This was done to account for the unequal stimulus-to-silence ratio of music and speech blocks, ensuring that activity was normalized to an unbiased baseline across different listening domains. Indeed, this local baseline normalization approach produced equal pre-stimulus baseline HFA values for music vs. speech vs. melodic speech in ERPs (e.g. Fig. 4e).

### Electrode localization

Electrodes were localized on each participant’s brain by co-registering the preoperative structural T1 MRI with postoperative tomography scans. Locations were superimposed onto a three-dimensional reconstruction of each patients’ cortical surface using a custom-written imaging pipeline (*73*). For electrode localization on a common atlas across patients, we used a nonlinear alignment procedure described in (*73*) to warp electrode locations from the patient’s native space to the cvs_avg36_inMNI152 template (*74*).

### Electrode selection

We first aligned continuous ECoG activity to the onset of musical phrases and viewed the peak of each electrode’s evoked response, averaged across all phrases, on a common brain map (fig. 1b). To screen for music-responsive electrodes, we applied a statistical test comparing the activity at each timepoint during sound presentation to activity during a silent period preceding each phrase (sign-rank p<0.05; Bonferroni correction for multiple timepoints and electrodes). Electrodes with a significantly higher magnitude of activity during sound presentation than baseline for a continuous window of at least 200ms were considered responsive and included in subsequent analyses (n = 224 music-responsive electrodes across 8 participants). The same statistical procedure was applied for evaluating speech-responsive sites, and we included the union of music and speech-responsive electrodes (n = 342) in all cross-domain analyses (Figures 3, 4, and 5). For an additional participant presented with synthesized harmonic/inharmonic tones, we identified sound-responsive electrodes using the same procedure as above (n = 7 electrodes).

### Single electrode sensitivity to contrasts in melodic features

To directly visualize whether responses at individual electrodes were sensitive to the pitch, pitch-change, or expectation of notes, we divided each feature’s distribution into two equal bins (median-split) and examined the corresponding cortical activity within each bin during the epoch from 100ms before to 400ms after the onset of notes (fig. 1c). To limit intrinsic correlations that exist between pitch and pitch-change (*r* = 0.14, p<0.001; figure S1), we excluded notes from the upper and lower quartile of the pitch distribution from this analysis only (we later use TRF modeling to overcome the issue of correlated features). Binned responses at each timepoint were compared using independent two-sample t-tests (p<0.001, Bonferroni corrected for timepoints, temporal threshold > 50ms). The above procedure was applied for every electrode to understand how activity was modulated by each melodic feature (fig. S2).

### Temporal receptive field (TRF) modeling of music evoked activity

To further quantify the extent to which different sources of melodic information were encoded in continuous activity at each electrode, we fit linear temporal receptive field (TRF) models. We discretized the pitch, pitch-change, and expectation of each phrase into *N* bins (see below), with each bin forming a unique row in the [*features x time*] stimulus matrix. We used binary predictors that were sparsely coded at note onsets to specify a given feature’s value (see fig. S3A). For pitch, we discretized values into 24 equally spaced (in log-Hz space) bins between the 5^th^ and 95^th^ percentile of the pitch distribution. Pitch values in the lowest and highest 5% of the distribution were placed into the first and last bins, respectively. By defining these percentile bounds, we prevent unstable estimates that can occur when extreme bins contain too few data points. The number of bins was chosen such that the lowest 12 pitch bins overlapped with the pitch distribution in speech (see below). We used the same approach to discretize both surprisal and uncertainty into 8 bins. For pitch-change, the distribution in music is naturally discretized into semitone bins. To avoid bins with little data points, we created distinct bins for each pitch-change between [-5, +5] semitones, and placed pitch-changes lower or higher than -5 and +5 semitones, respectively, in two additional bins. In addition to modeling neural responses to the three melodic features, the full TRF model also included the auditory spectrogram (a peripheral stimulus representation), which we extracted using the NSL toolbox (*75*) and downsampled to 25 logarithmically spaced bands spanning 6 octaves between 0.08 – 8kHz. We also included two binary predictors specifying the onset location of phrases and notes, respectively. These temporal landmark features allow us to statistically control for the contribution of onset-from-silence responses (*76*) and the presence or absence of a note when evaluating the contribution of specific melodic information. Finally, we included a sparse predictor at note onsets that specified the number of times a note with the same pitch had consecutively occurred previously within the phrase. This “note repeat” predictor controls for the effects of stimulus specific adaptation that may be confounded with low expectation contexts. Before fitting, predictors were scaled between 0 and 1 by dividing them by the magnitude of the maximum value in each bin. This ensured that all estimated beta values were scale free and comparable across predictors, with beta magnitude being an index for the contribution of a predictor to model performance.

Neural activity at each timepoint *HFA(t)* was modeled as a weighted linear combination of features of the stimulus *X* within a window spanning *t–400ms* to *t+100ms*. For each feature *f*, this resulted in a set of weights, *b*_*1…, d*_, with *d* = 50 for a sampling frequency of 100 Hz across the 500 ms window (fig. 2B and fig. S3).

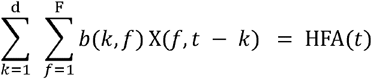

Models were estimated separately for each electrode using L_2_ regularization (ridge regression) and fivefold cross-validation. For each cross-validation fold, we trained models on 80% of the data and evaluated them on the held out 20%. The regularization parameter was estimated using a 10-way bootstrap procedure within each training fold before a final value was chosen as the average optimal value across folds. Model performance was evaluated as the Pearson’s correlation between actual and predicted brain responses. These correlations were squared to obtain the *R*^*2*^, a measure of the portion of variance in neural activity explained by the model (fig. 2a and supplementary fig. S3)

### Noise ceiling

To evaluate how well TRF models performed relative to an upper limit on explainable variability, we computed the noise ceiling for each electrode using the adjusted split-half correlation approach (*36, 37*) applied to a subset of musical phrases that were repeated 11 times to participants. First, we randomly split trials into 2 groups and computed the R^2^ value between the averages of the two groups. We repeated this process many times to obtain an average of the distribution of values, which we inserted into the equation:

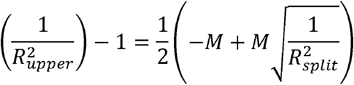

Where *M* represents the number of stimulus repetitions. We divided true TRF *R*^*2*^ values by these noise-ceiling estimates to obtain the distribution of noise-corrected *R*^*2*^ values (Fig. 2a).

### Variance partitioning of TRF models

To estimate the contribution of a specific feature to the full TRF models (fig. 2b-c), we computed its unique variance explained (Δ*R*^*2*^). For a given feature of interest *G*, we fit a reduced TRF model that predicted neural activity using all features except *G*. We then evaluated the unique variance explained by *G* as the difference in *R*^*2*^ between the full and the reduced models:

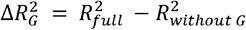

The unique variance explained by expectation was estimated as the combined contribution of both surprisal and uncertainty. The significance of each feature’s unique variance contribution was calculated using a permutation test. Specifically, we shuffled the rows of the predictor matrix corresponding to a feature of interest, leaving all other rows (corresponding to other features that were not being tested) intact. We then fit a TRF model to the permuted predictor matrix. Repeating this procedure 200 times, we arrived at a null distribution. True Δ*R*^*2*^ values that fell above the 95^th^ percentile of these permuted Δ*R*^*2*^ values were considered significant.

### Independent encoding of melodic pitch, pitch-change, and expectation

To determine whether the three melodic features were encoded by overlapping or independent neural populations, we directly compared encoding using linear mixed-effects models. Specifically, for each of the 3 electrode subsets that significantly encoded pitch, pitch-change, and expectation respectively, we modelled the response variable as the encoding (Δ*R*^*2*^) of a given feature. We included fixed effects terms for the encoding of the other two features. We also included random effects for the intercept and slope grouped by participant. Inclusion of these random effects terms ensured that results were not driven by unequal representation of a given participant in pooled data.

### Anatomical organization of melodic encoding

To examine whether the encoding of pitch, pitch-change, and expectation was spatially organized along the STG (fig. 2d), we extracted electrode locations along the posterior-anterior axis of the brain and normalized each participant’s locations to the point at which the central sulcus meets the sylvian fissure. Normalizing to a common anatomical landmark avoids warping individual’s electrode locations to a common brain, better preserving the relative spatial differences between encoding of each feature. Within each hemisphere, we assessed whether the distribution of electrodes that encoded each feature was significantly different in its posterior-to-anterior location using linear mixed-effects models with a randomized block design in which blocks correspond to participants.

### Melodic feature tuning

For electrodes that significantly encoded a melodic feature (based on TRF Δ*R*^*2*^), we sought to further characterize their tuning patterns – that is, the format by which information was encoded. In addition to inspecting corresponding TRF weights (Fig. 2e), we examined tuning patterns in the raw HFA using the same bin edges as that used in constructing TRF stimulus matrices. To produce time-collapsed tuning matrices (fig. 2f-h), we estimated each electrode’s peak encoding latency *K* relative to the onset times of notes in the stimulus. This estimate was based on the lag at which the magnitude of TRF weights was maximal. For every musical note onset at time *t*, we then binned the HFA at time *t + K*. Responses in each bin were then averaged, and tuning across bins was normalized by removing the mean and dividing by unit variance.

To further quantify tuning across pitch-encoding electrodes, we calculated the F0 corresponding to the peak-HFA in tuning curves for each electrode (Fig. 2f, right). To examine whether tuning to pitch could be accounted for by correlated variables, we examined whether tuning existed independent of (i) the spectral profile of notes, the centroid of which was strongly correlated with note F0 (r = 0.75; Fig. S4), or (ii) the spectral modulation profile, the centroid of which was correlated with, but also contained substantial independent variation from, F0 (r = -0.4; Fig. S5a-b). To dissociate between spectral profile and F0, we examined pairs of note-clusters that spanned comparable pitch-ranges but differed in their spectral profiles (Fig. S4b-c). This was achieved by grouping notes by instrument or by manually selecting clusters of notes with anti-correlated spectral profiles but with similar F0 ranges. For every electrode, we characterized F0 tuning separately within each cluster, and correlated the two corresponding tuning curves (Fig. S4e). To test whether F0 tuning could be explained by tuning to rates of spectral modulation, we first extracted the spectral modulation profile of notes using the nsltoolbox (*75*), using 16 bins between 0.25 - 4 octaves per cycle. We then included the 16 spectral modulation bins as predictor variables in TRF models (along with other melodic and acoustic predictors). To determine whether the observed F0 tuning was independent of spectral modulation tuning, we assessed (i) whether replacing F0 TRF predictors with spectral modulations yielded lower overall model R^2^ values (Fig. S5c) and (ii) whether pitch still explained unique variance in activity when spectral modulation variables were already included in the model (Fig. S5d).

For pitch-change encoding electrodes, we used k-means clustering to group electrodes with similar tuning profiles across the range of pitch-changes (Fig. 2g, right). We chose k=3 as this produced clearly distinct and interpretable clusters. We sought to determine whether the encoding of pitch-change specifically reflected a comparison of F0 between two adjacent notes or a comparison of their individual frequency components. We took two complementary approaches to address this question. Firstly, we asked whether tuning to pitch-change was dependent on whether high-order frequency components (above the F0) provided a reliable cue to pitch-change direction (ascending or descending). We first classified every pitch-change in the music stimulus based on whether components (excluding F0) provided an unambiguous versus ambiguous cue to the direction of pitch-change (Fig. S6a-b). To classify a pitch-change as either ambiguous or unambiguous, for every harmonic component of a given order, we determined whether the most proximate component (in log-Hz space) within the next note had the same or different order. Pitch-changes in which at least 50% of all nearest-components were of a different order were classified as ambiguous. Consistent with prior research, we found ambiguity primarily arising for larger interval-magnitudes that were greater than 3 semitones (*38*). We then asked whether the degree of response modulation to ascending versus descending pitch changes differed across ambiguous versus unambiguous conditions (Fig. S7c-d). In a second approach, we analyzed neural data from one participant presented with synthesized melodies comprised of either harmonic or inharmonic tones in addition to music. Due to limited electrode coverage over auditory regions in this participant, we found relatively few sound-responsive electrodes across both music and harmonic tones (n = 7, see electrode selection procedure above). However, we identified one electrode whose responses were significantly modulated by the direction of pitch-change in melody (independent two-sampled t-test, p<0.05, temporal-threshold > 50ms). Using the same approach, we analyzed whether responses at the same electrode were also modulated by pitch-change direction during presentation of harmonic or inharmonic tones (Fig. S6c, top panels). To understand whether effects were driven by small vs. large pitch-changes, we grouped time-averaged responses into two bins corresponding to small (≤ 3 semitones) vs. large (> 3 semitones) pitch-change magnitudes and quantified response differences between ascending vs. descending changes separately in each bin, using a bootstrapping procedure to generate 95% CIs for statistical inference (Fig. S6c, bottom panels). Within this participant, we also asked whether pitch or expectation were encoded across music, harmonic and inharmonic conditions. We identified one electrode whose responses were significantly modulated by F0 across both music and harmonic-tone conditions (Pearson’s correlation between F0 and HFA r > 0.2 for both conditions, p < 0.001) but not the inharmonic condition (r = 0.04, p > 0.05). We found no electrodes significantly modulated by expectation in any condition (independent two-sample t-test on median-split responses, all p>0.05).

For expectation encoding electrodes, we tested for monotonic encoding by computing the rank order correlation between the response to each note and its corresponding expectation value (Fig. 2h, right). We also plotted ERPs by dividing the distribution of expectation into 4 equal-spaced bins and grouping corresponding neural responses (Fig. S9a).

### Domain selectivity

To characterize the extent to which activity at each electrode was stronger for music or speech (fig. 3a), we derived a domain selectivity index (SI) ranging from -1 (speech selective) to +1 (music selective). For each electrode, we concatenated activity evoked during the first second of music and speech tokens, respectively, into two vectors, which we then compared using an independent two-sample t-test. We used the resulting test-statistic as a proxy for domain-selectivity, normalizing values so that the largest magnitude across all electrodes was equal to 1. To determine whether music-selectivity anatomically clustered (fig. 3c), we extracted electrode locations along the posterior-anterior axis of the brain and normalized each participant’s locations to the point at which the central sulcus meets the sylvian fissure. Within each hemisphere, we then compared the location of music-selective electrodes (SI > 0.2) with that of all other electrodes (SI < 0.2) using linear mixed-effects models with a randomized block (where blocks correspond to participants). We further conducted within-participant comparisons of the location of music-selective versus nonselective electrodes using Bonferroni-corrected ranksum tests. To ensure results were not due to our specific choice of a cutoff SI = 0.2, we repeated these analyses using SI = 0.33 as the cutoff and verified that results remained unchanged. Specifically, linear-mixed effects analyses confirmed that music-selective electrodes did not anatomically differ from other electrodes in both left (t = 1.15, p = 0.25) and right (t = 0.23, p = 0.82) hemispheres, and all within-subject comparisons between the two distributions were insignificant (p > 0.1, ranksum tests).

### Relationship between feature encoding and selectivity

To determine whether music-selectivity was explained by the encoding of pitch, pitch-change, or expectation, we computed the partial correlation between the Δ*R*^*2*^ explained by each feature and the SI across all electrodes with SI>0 (Fig. 3e, Fig. S10). For a given feature, partial correlations controlled for the Δ*R*^*2*^ explained by the other two features. Partial correlations also controlled for the magnitude of the mean music response on each electrode (electrodes with stronger music responses were more likely to be music-selective and have greater Δ*R*^*2*^ values by default, leading to spurious correlations). Significance was evaluated by randomly permuting the SI and Δ*R*^*2*^ values before correlating permuted variables. This procedure was repeated 10,000 times to define a null distribution. Cases in which the true correlation exceeded the 95% of the null distribution were considered significant. Similarly, to determine whether speech-selectivity was explained by the encoding of relevant features of speech (fig. 3g, see below), we applied the same procedure as above, evaluating partial correlations between Δ*R*^*2*^ and SI across all electrodes with SI<0.

### TRF modeling of speech

Speech-evoked activity at each electrode was modeled using the same TRF approach that was used to model music activity. We specifically modeled features of speech that were analogous to melodic pitch, pitch-change, and expectation. To code pitch in the stimulus matrix, we discretized the continuous pitch contour of sentences into 12 equally spaced bins. To extract pitch-change, we computed the difference between the median pitch of the current and prior syllable, discretized values into semitone-spaced bins, and coded each interval as a separate row in the stimulus matrix at the onset of syllable nuclei. To consider sequence statistics in speech that are comparable to melodic expectation, we extracted and modeled both phoneme surprisal and cohort entropy, as defined by (*42*). These values are mathematically equivalent to those calculated earlier for music, with phonemes in place of notes. Specifically, phoneme surprisal is the inverse of the conditional probability of each phoneme give the preceding phonemes in a word:

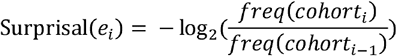

where *cohort*_*i*_ is the cohort of words at phoneme with position *i*, and *freq(c)* is the summed frequency of all words in cohort *c*. Cohort entropy is the Shannon entropy of the cohort at each phoneme, given by:

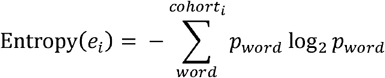

Where *p*_*word*_ is the relative frequency count of the given word within a language corpus (see *42* for details). We specified these values in the stimulus matrix at the onset of each phoneme and discretized both variables into 8 bins. To control for the contribution of extraneous speech features, we included the spectrogram and two binary predictors specifying sentence and syllable-nuclei onset locations. All other aspects of the TRF fitting procedure and estimation of unique variances were identical to the modeling approach used for music.

### Analysis of melodic speech

To evaluate the extent to which each melodic speech token elicited the percept of melody (Fig. 4d), we ran a behavioral test on an independent set of listeners (N=11). Subjects rated the extent to which each token elicited the percept of melody on a continuous scale of 0 – 10, and were explicitly instructed to use regular speech and song as perceptual anchors that correspond to ratings of 0 and 10 respectively.

To assess the extent to which electrodes responded more strongly to melodic over regular speech (fig. 4f), we computed the normalized difference in evoked activity at each electrode as:

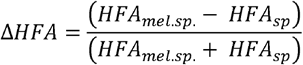

We evaluated the ΔHFA for electrodes within three different groups: speech-selective (SI<-0.2), shared (−0.2<SI<0.2), and music-selective (SI>0.2). Within each bin, we used a one-sample t-test to assess whether the mean ΔHFA was significantly greater than zero, which would indicate enhanced responses to melodic versus regular speech. To evaluate whether responses to melodic speech tokens scaled with the degree to which they were perceived melodically, we correlated the average ratings across listeners with the corresponding peak HFA elicited by each token (fig. 4g). We computed this correlation for all electrodes and grouped values into the same three SI-based bins as above (fig. 4h). Within each bin, we used one-sample t-tests to evaluate whether mean correlations were significantly different from zero.

We examined feature tuning in melodic speech by first extracting pitch, pitch-change, and expectation (again using melodyRNN) from every token. We divided each feature distribution into two bins (using the median-split) and inspected corresponding neural responses in each bin. Because melodic speech tokens with female speakers were rated as more melodic than those with male speakers (Wilcoxon rank-sum: p = 6.65×10^−5^, z = -4), to avoid speaker-identity driven effects, we binned neural responses separately for male and female subsets of the data, before averaging across gender. Additionally, we did this separately for tokens that were perceived as more versus less melodic (using the median split of perceptual ratings). The variance in expectation for tokens on either side of this median split did not differ (t = 0.57, p = 0.56). For a given feature and electrode, we compared neural responses in the two bins using independent-sample t-tests and used the resulting test-statistic as a proxy for the strength of tuning to that feature. As was done earlier with music, we used partial correlations to predict SI from tuning to melodic features, evaluating significance using permutation-based null distributions (fig. 4j).

### Cross-domain comparison of pitch and pitch-change encoding

To compare encoding across domains, across all electrodes that encoded a given melodic feature, we computed the linear correlation between the Δ*R*^*2*^ values explained by pitch or pitch-change in music versus speech (fig. 5d). To compare feature tuning curves between music and speech (fig. 5e), we confined our analysis to the range of overlapping F0 and interval values across domains (fig. 5c). For F0, we focused on the range from [86 Hz, 252 Hz], corresponding to the 5^th^ – 49^th^ percentile and the 2.5^th^ – 97.5^th^ percentile of the pitch distributions in music and speech respectively. For interval, we focused on the set of 9 intervals from [-4, +4] semitones, corresponding to the 9^th^ – 91^st^ percentile and the 3.5^th^ – 98^th^ percentile of the pitch-change distributions in music and speech respectively. To characterize pitch tuning, we grouped neural responses to notes (for music) or to voiced portions of an utterance (for speech) into 12 log-Hz spaced bins. Relative to the onset time of a pitch event in the stimulus, we estimated each electrode’s peak encoding latency *K* based on the lag at which the corresponding pitch-TRF weights were maximal. For every musical note or voiced portion of speech occurring at time *t*, we binned the HFA at time *t + K*. Next, we averaged responses in each bin and centered the data for each electrode (removed the mean and divided by unit variance) to clearly visualize modulations in mean neural activity across the range of pitch. We used the same approach to compare pitch-change tuning across domains. We quantified the similarity in neural codes across music and speech by computing the linear correlation between tuning curves across all electrodes.

### Cross-domain decoding of pitch and pitch-change

To quantify the extent to which neural codes for pitch and pitch-change generalized across domains (fig. 5f), we used linear discriminant analysis (see (*77, 78*) for details on the decoding procedure). We trained classifiers to decode either pitch or the contour of pitch-change from cortical responses across all electrodes that significantly encoded a given feature in both domains. As with tuning matrices above, we only used the HFA at the timepoint corresponding to peak TRF encoding relative to the stimulus for decoding. For pitch, we discretized the distribution into 4 classes (chance accuracy= 25%). For pitch-change, we classified activity for descending versus ascending changes (chance accuracy = 50%). For the purposes of evaluating generalization of pitch-change, we chose to classify contour (ascending vs. descending) rather than specific interval information, as intervals are not well defined in speech. To evaluate decoding accuracy within-domain, we used 10-fold cross validation to maintain independent test and train data sets. For decoding across-domain (e.g., train on music, test on speech), we utilized all the data in each domain to train and test models. Before decoding, we equalized the number of observations of each class (observations per class for pitch: *N = 355* for music, *N = 6598* for speech; observations per class for pitch-change: *N = 1243* for music, *N = 689* for speech) and centered the data in each domain to have mean = 0. Statistical inference was performed by bootstrapping data and re-applying the entire decoding analysis pipeline on each run (N runs = 200).

## Acknowledgements

We thank all members of the Chang Lab for useful feedback and discussion. We thank Ben Speidel for help with anatomical localization of electrodes.

## Author contributions

Conceptualization: NS, MKL, EFC; Methodology: NS, MKL, FT; Software: NS; Formal Analysis: NS; Investigation: NS, MKL, EFC; Data Curation: NS; Writing – original draft: NS; Writing – review and editing: NS, MKL, FT, EFC; Visualization: NS; Supervision: EFC, MKL, FT; Funding Acquisition: EFC.

## Data and materials availability

Data will be available upon reasonable request to the corresponding author.

## Supplementary Materials

Figs. S1 to S10

Tables S1 to S2

Audio S1 to S3

