## supplementary figures 1 - 10. Tables 1 -2 for "Encoding of melody in the human auditory cortex"

**This PDF file includes:**

Figs. S1 to S10

Tables S1 to S2

Captions for Audio S1 to S3

**Other Supplementary Materials for this manuscript include the following:**

Audio S1 to S3

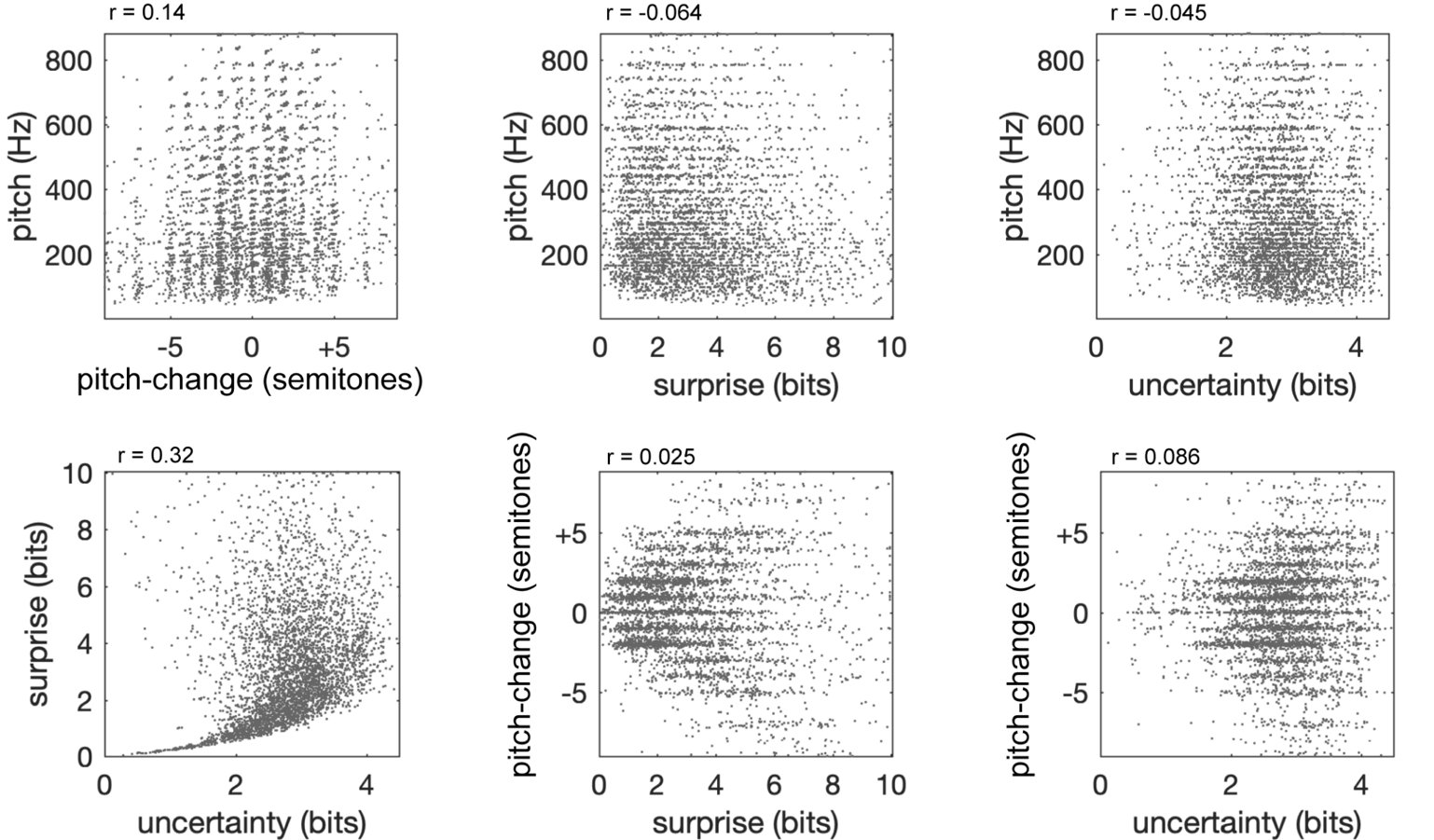

**Figure S1. Correlations amongst melodic features.** Gray markers represent individual notes in the music stimulus set. Each panel illustrates the correlation between two melodic features. For expectation, correlations are computed for both surprisal and uncertainty (though we focused on surprisal; see Fig. S2 below). Linear correlation coefficients (Pearson’s *r*) indicate the strength of each correlation.

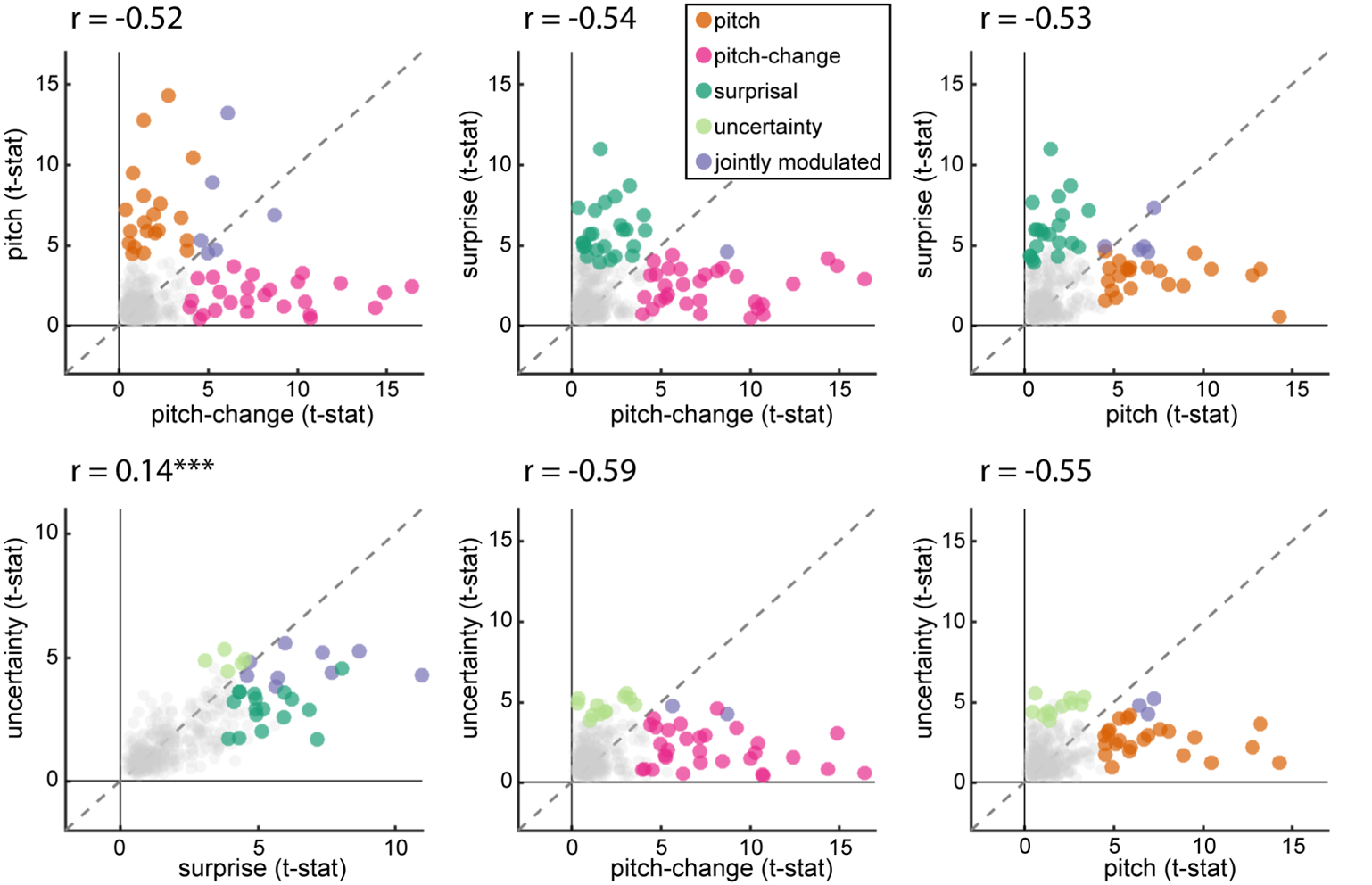

**Figure S2. Response modulation due to melodic features.** Each feature’s distribution is divided into two bins (using the median split). For every electrode, neural responses in the two bins are compared using independent two-sample t-tests, and the resulting test-statistic is used as a proxy for the strength of modulation. Pitch, pitch-change, and expectation modulate neural responses on distinct subsets of electrodes. Surprisal and uncertainty modulate responses at the same electrodes, with surprisal exerting a larger modulatory effect than uncertainty. Electrodes at which responses were significantly modulated by a given melodic feature (p<0.05) are indicated by marker colors. Asterisks represent significance level for linear positive correlations of values on x and y axes (***p<0.001).

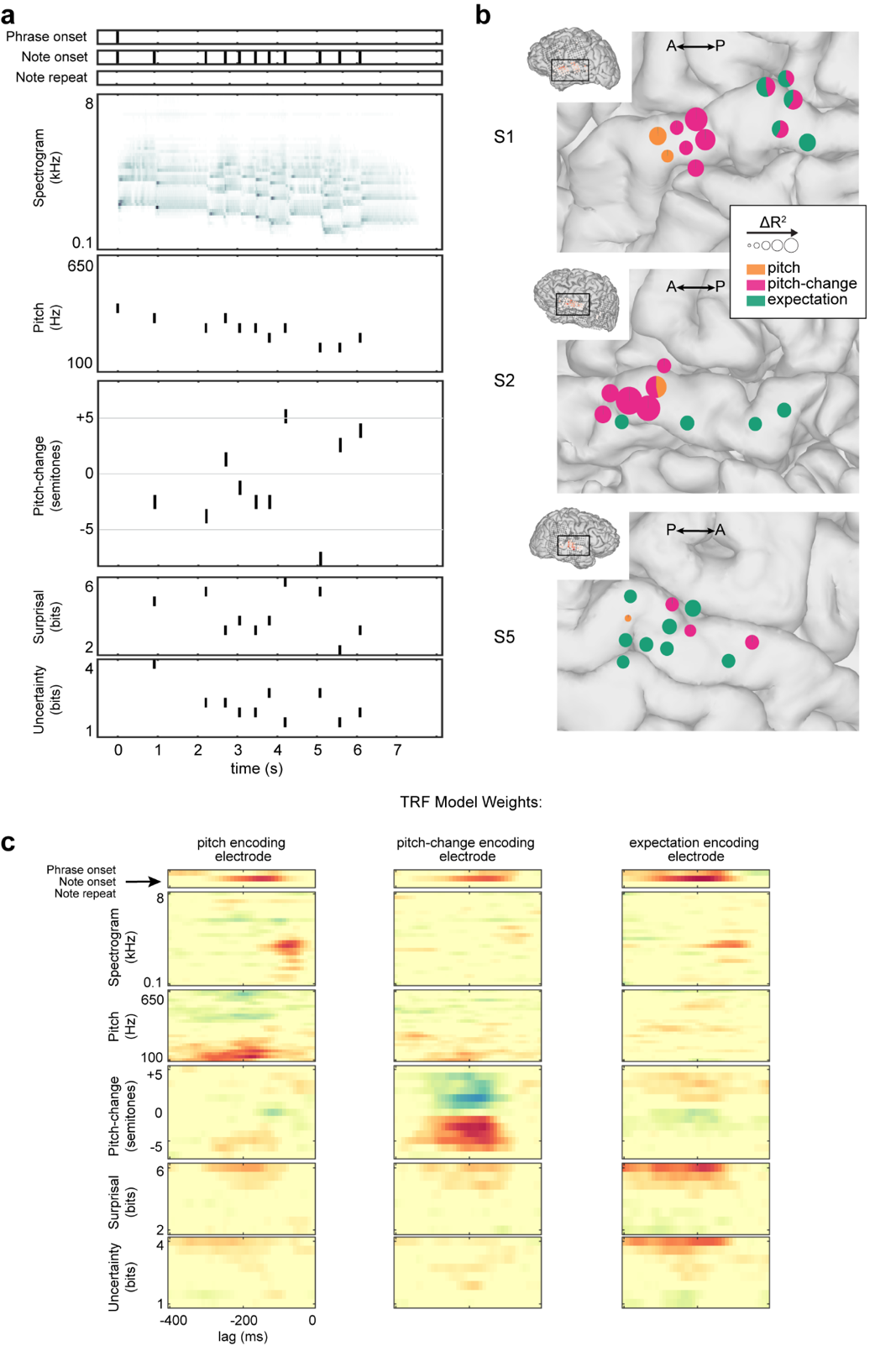

**Figure S3. TRF modeling of melody.** (A) Stimulus design matrix used for modeling continuous neural responses to music. (B) Unique variance of melodic features plotted on cortical reconstructions of three participants (2 LH, 1 RH). (C) TRF model weights for example electrodes tuned to different melodic features.

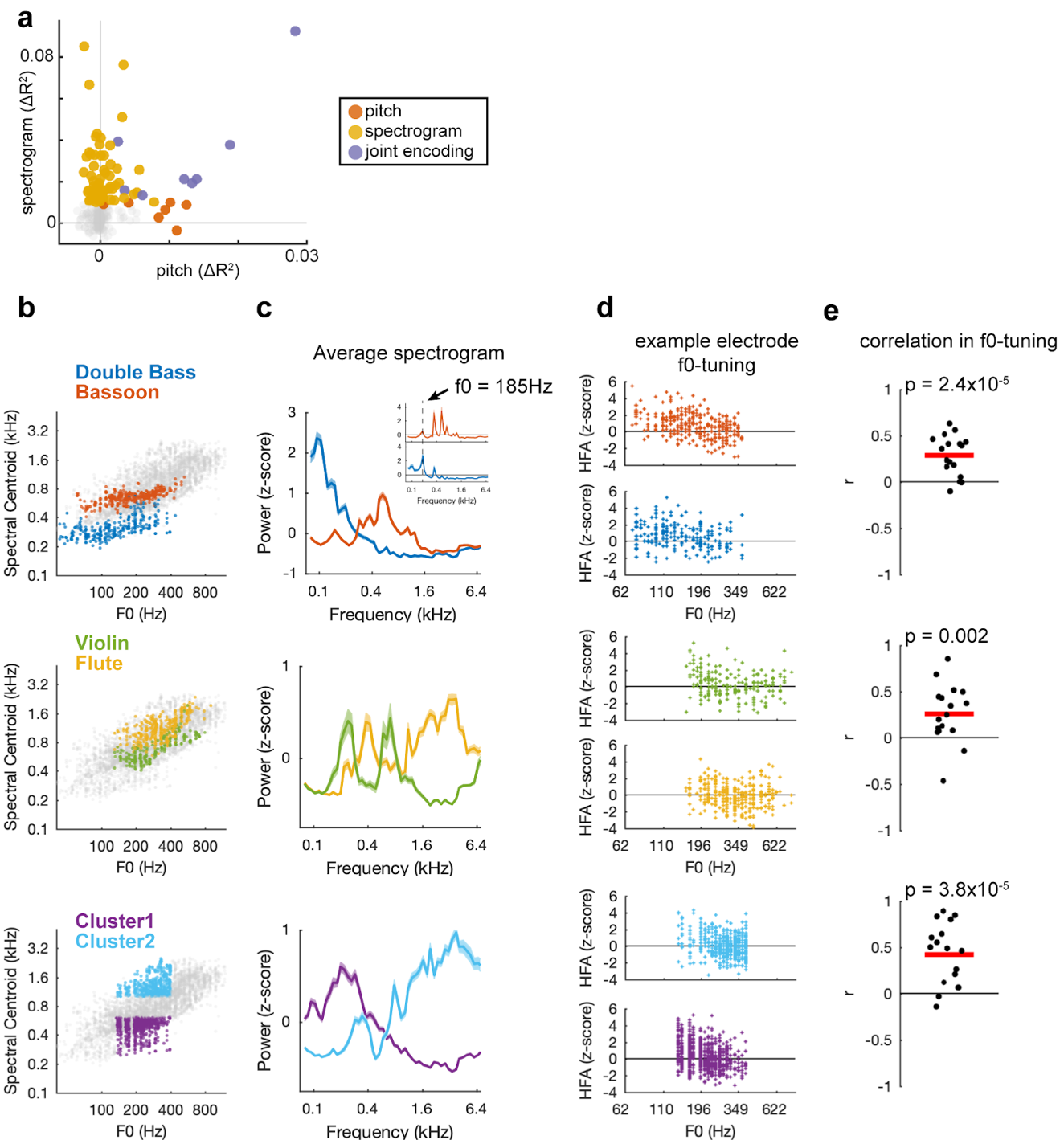

**Figure S4. Pitch-encoding is independent of the spectral profile of notes.** (A) Pitch and spectrogram features explain variance in activity at partially overlapping electrode subsets. (B) Scatter plots of notes’ F0 versus spectral centroid. Although highly correlated across the entire stimulus (r=0.75), dissociation of F0 and general spectral information can be achieved by comparing stimulus subsets (colored markers) with highly overlapping F0 but distinct spectral profiles. (C) Average spectral profiles of note-clusters being compared. Top inset: average profile for a fixed F0 value. (D) Markers show the neural response to individual notes for an example electrode. Similar pitch tuning exists across spectrally dissimilar note-clusters. (E) Across electrodes, a positive correlation exists between the F0-tuning curves to spectrally dissimilar notes (t-test: t-stat >3.4, p< 0.001). Each marker represents a pitch-encoding electrode.

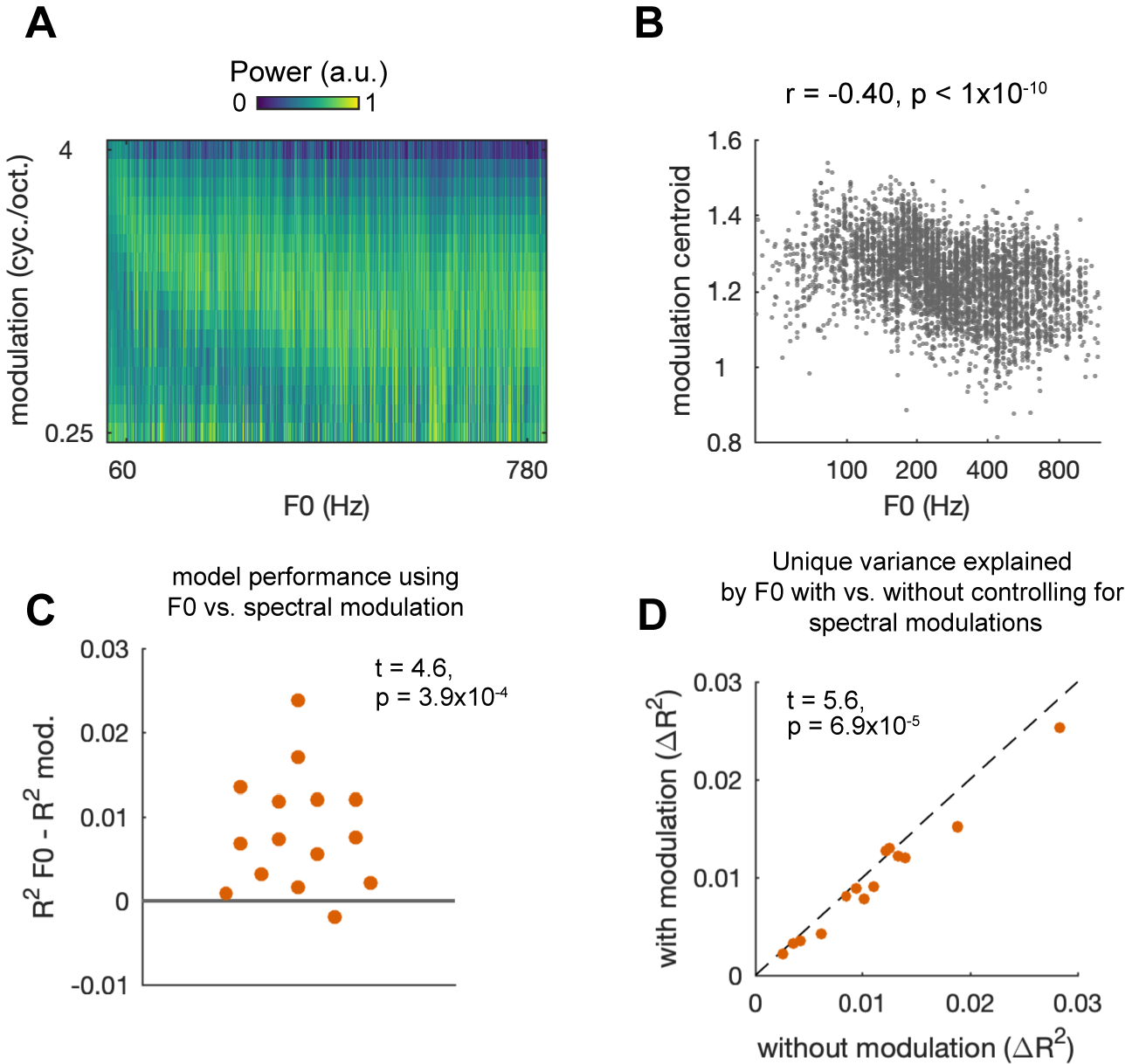

**Figure S5. Pitch-encoding is independent of tuning to spectral modulation rates.** (A) Modulation power (y-axis) across all notes sorted in order ascending F0. (B) Each marker represents a note. Scatter plot shows a moderate negative relationship between pitch and spectral modulation centroid across all notes, indicating that notes with lower pitch have higher modulation rates (r = -0.4). Note however that there exists substantial variability in modulation rates at a given f0. (C) Replacing F0 predictors with spectral modulation predictors in TRF models yields significantly lower overall performance. Y-axis plots the difference in model R^2^ when using F0 versus spectral modulations as a proxy for pitch. X-axis does not represent any variable. (D) Relative to a TRF model that does not control for spectral modulations (x-axis), pitch still explains *∆R^2^* (though significantly less) when controlling for spectral modulations in TRF models (y-axis), suggesting that the neural variance explained by pitch is largely independent from any variance that may be attributed to spectral modulation.

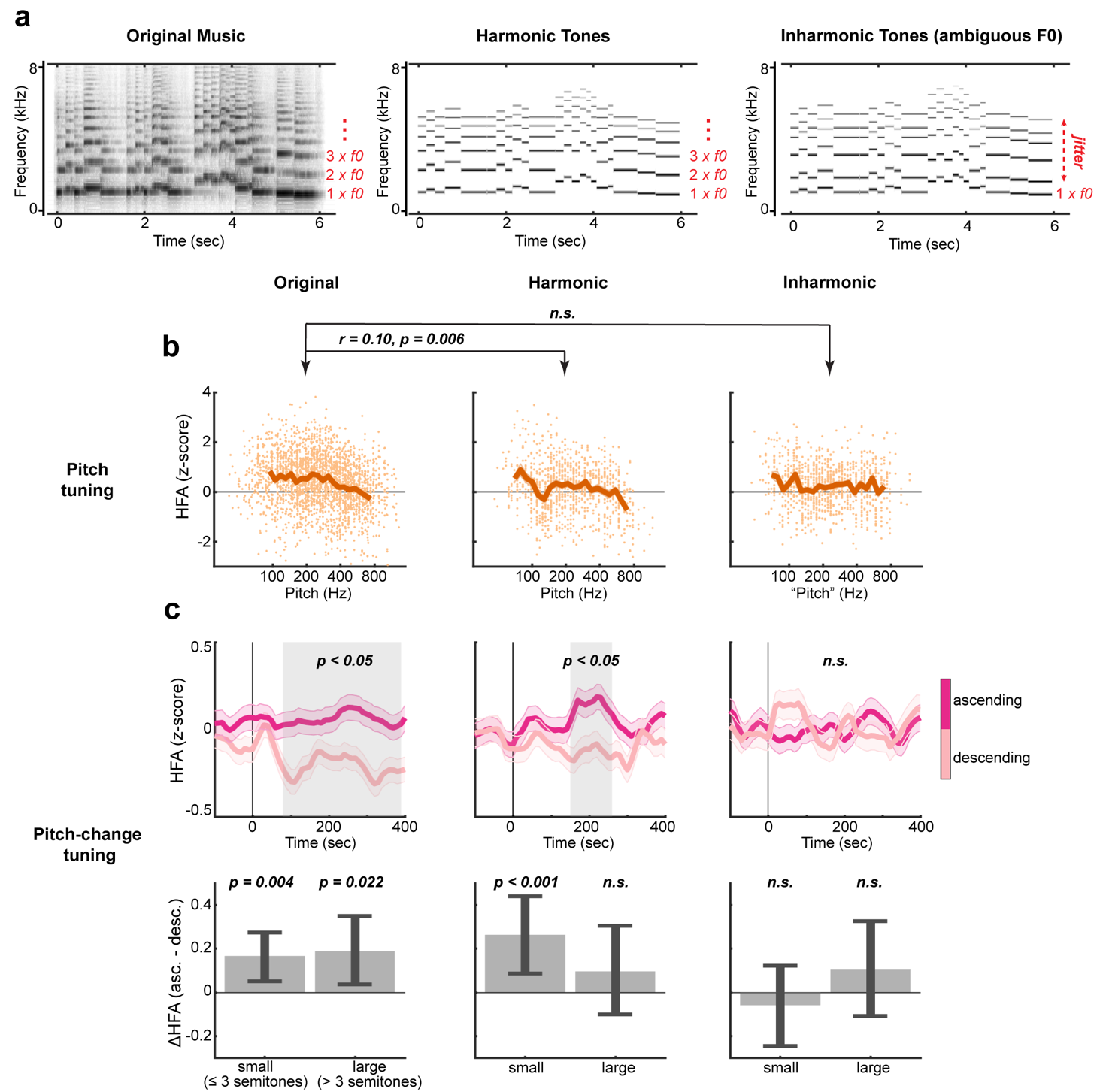

**Figure S6. Tuning to pitch and pitch-change across original music, harmonic, and inharmonic complex tones.** (A) Spectrograms for an example melody in its original form (left) or synthesized as harmonic (middle) or inharmonic (right) tones. Inharmonicity was created by pseudo-randomly jittering harmonic components above the F0 (see methods). (B) Tuning to pitch at one electrode across the three listening conditions. Pitch tuning in original music was positively correlated with pitch tuning in harmonic (r = 0.1, p = 0.006) but not inharmonic (r = 0.059, p = 0.11) melodies. (C) At a different electrode, responses are significantly tuned to ascending pitch-changes in both original music and harmonic melody, but not inharmonic melody. Grey shaded regions indicate timepoints at which response differences to ascending vs. descending contour were significant (t-test, p<0.05). Bottom: Average HFA modulation due to pitch-change (ascending – descending) separated by interval magnitude (small vs. large). Error bars indicate 95-CIs derived from bootstrapping.

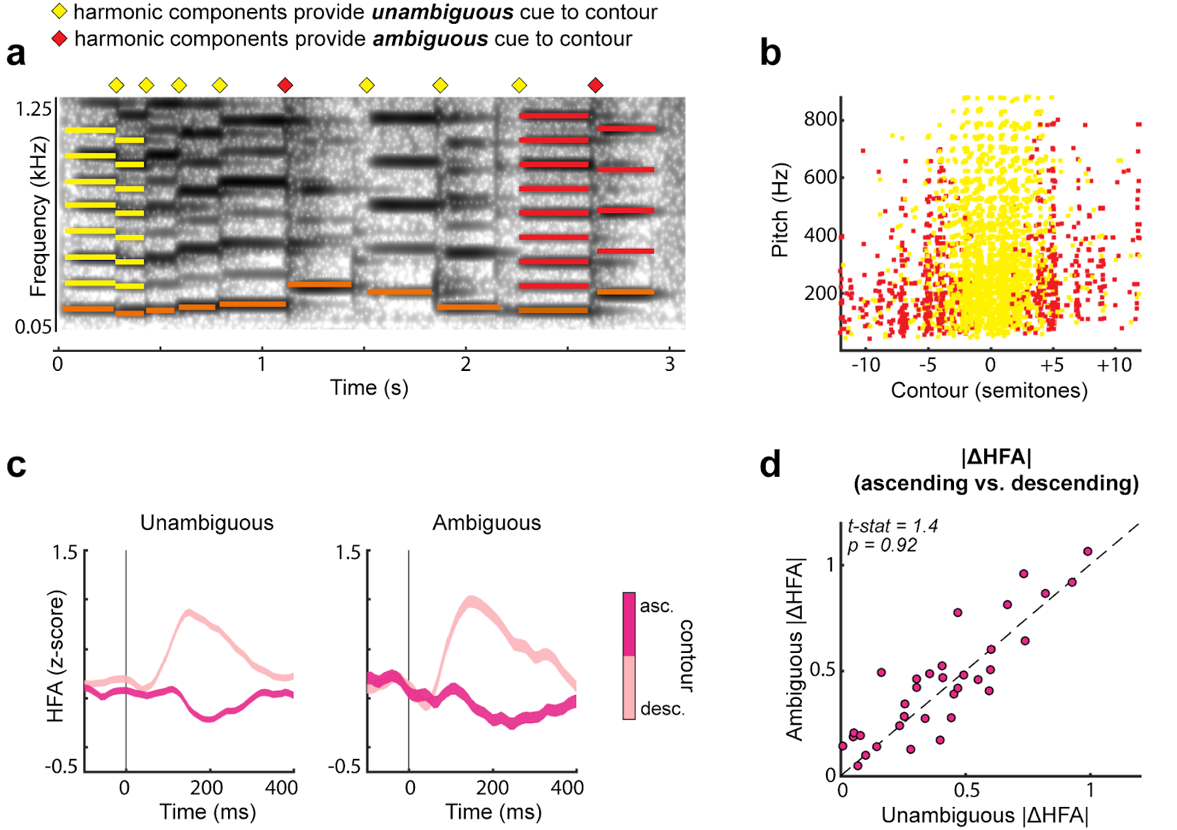

**Figure S7. Tuning to the direction of pitch-change is independent of whether harmonic components provide reliable cues.** (A) Spectrogram for an example melody. Colored diamonds above indicate pitch-changes in which harmonic components provide an unambiguous (yellow) or ambiguous (red) cue to the melodic contour. For instance, between the first two notes, harmonic components are close in frequency (yellow lines) and thus provide an unambiguous cue to contour. Such a correspondence between components is not present in the pitch-change between the last two notes (red lines), leaving only the f0 as a reliable cue. (B) Consistent with prior work, ambiguity primarily occurs at large interval sizes (≥ ±4 semitones). Across the stimulus, 22.2% of all pitch-changes contained ambiguity. (C) Response traces for an example electrode show similar selective responses to descending pitch-change for both unambiguous (left) and ambiguous (right) conditions. (D) The magnitude of response modulation did not differ between unambiguous (x-axis) and ambiguous (y-axis) instances across all pitch-change encoding electrodes.

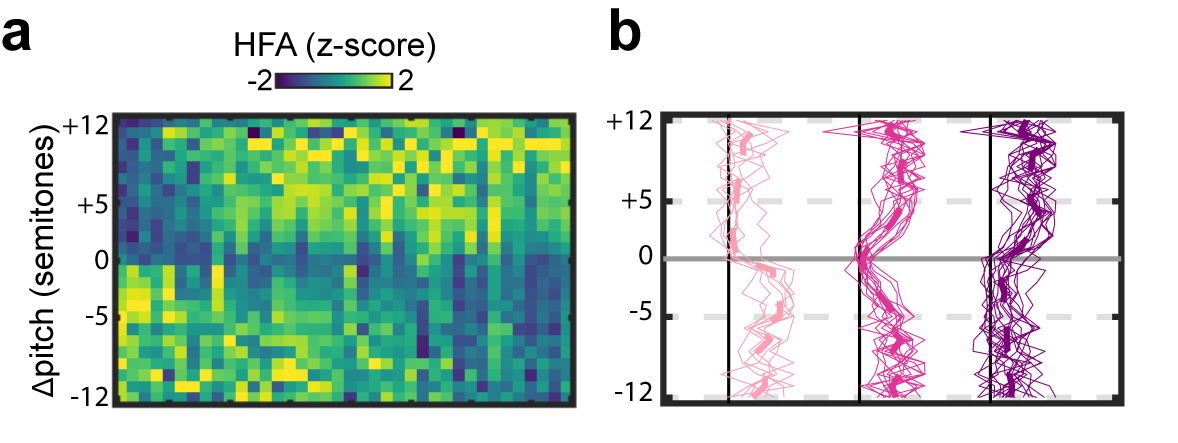

**Figure S8. Tuning to pitch-change across small and large interval sizes in the musical stimulus.** (A) Pitch-change tuning curves for all pitch-change encoding electrodes across the range from -12 to +12 semitones. Relatively few data points exist in the stimulus beyond interval magnitudes of 5 semitones (see fig. 5c). (B) Clustering (k-means, k = 3) of tuning curves.

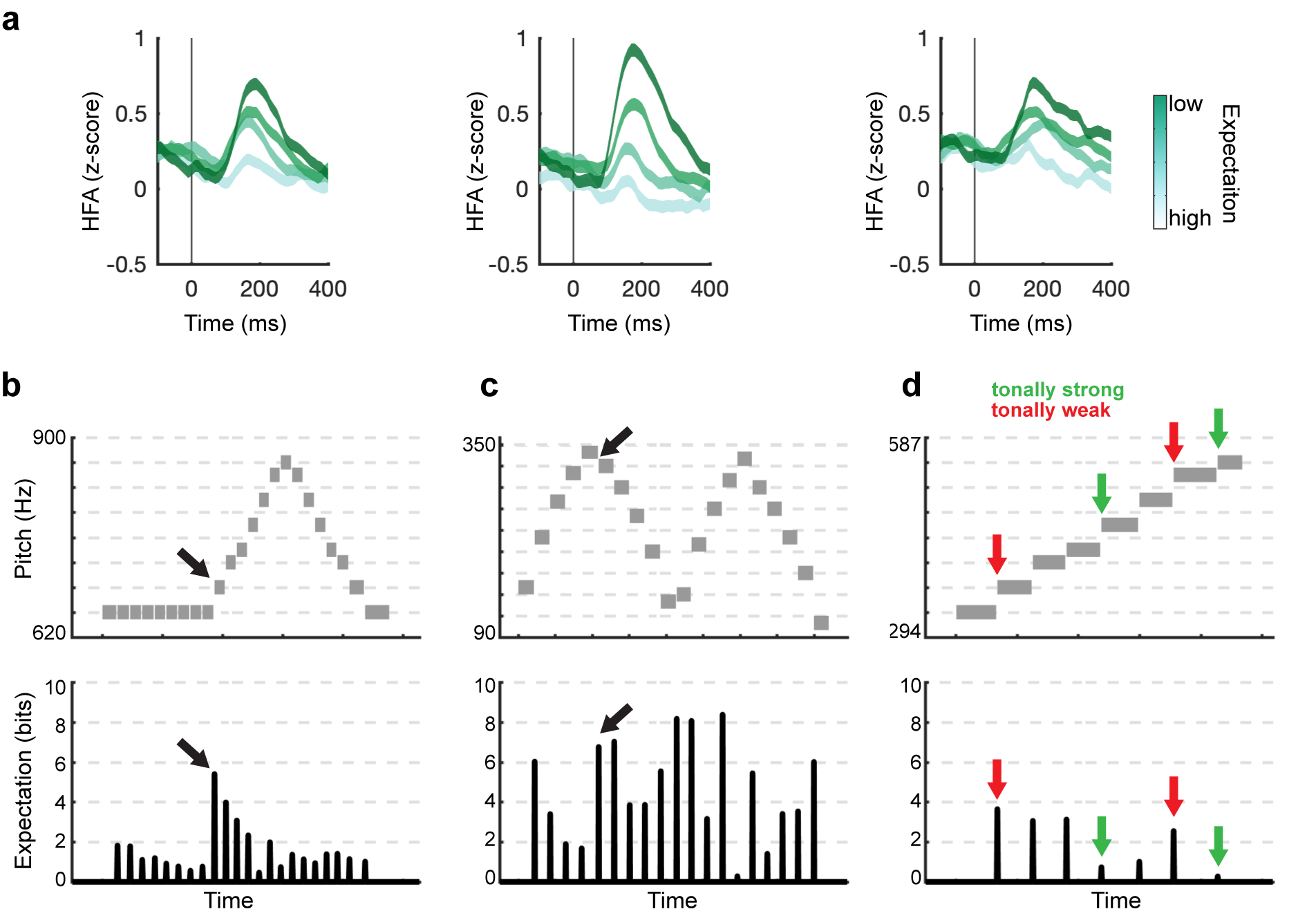

**Figure S9. Expectation reflects violation of sequence patterns and structural rules of Western tonal music.** (A) Note-evoked responses for three expectation-encoding electrodes, showing progressively larger responses to more unexpected notes. (B-D) To demonstrate the relationship between characteristics of melody and expectation, each panel shows the pitch track (top) and corresponding expectation values (bottom) for three example phrases. Black arrows in panels B-C indicate notes violating acoustic sequence patterns of pitch and contour. In panel D, the example phrase consists of an ascending major scale and demonstrates higher-order effects of tonality. Specifically, tonally strong pitches within the musical key (green arrows, referred to as the tonic and dominant) are more expected, while tonally weak pitches (red arrows) are less expected.

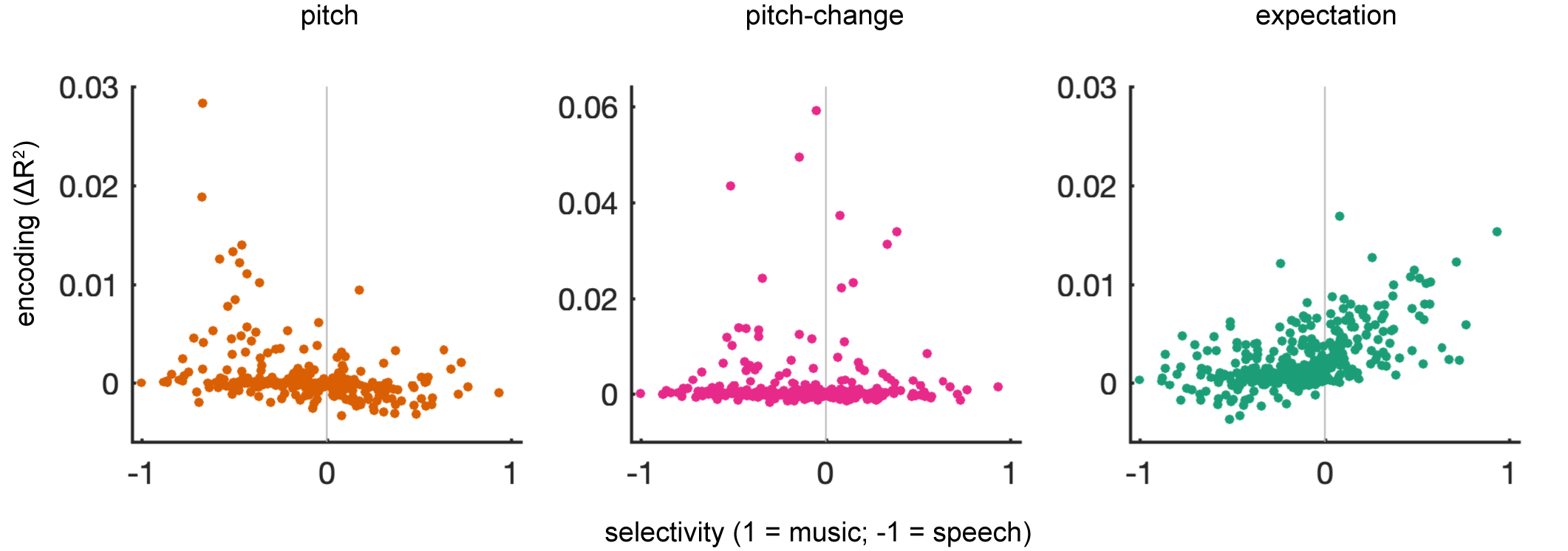

**Figure S10. Relationship between selectivity and melodic feature encoding.** Scatter plots show the unique variance explained by each feature of melody against selectivity. Encoding of expectation is proportional to music selectivity.

| **Subject** | **Hem.** | **Age** | **Sex** | **Handedness** | **Years of music training** | **L1** | **Epilepsy Focus** |
| --- | --- | --- | --- | --- | --- | --- | --- |
| EC214 | L | 39 | F | L | 0 | Spanish | Left hippocampus and anterior temporal lobe |
| EC222 | L | 47 | F | R | 7-8 | English | Left mid-posterior lateral and basal temporal lobe |
| EC225 | L | 31 | M | R | 0 | Spanish | Left occipital and anterior temporal lobe |
| EC228 | L | 46 | F | R | 0 | English | Left posterior basal temporal lobe |
| EC235 | R | 36 | F | R | 1 | English | Right anterior temporal lobe |
| EC242 | R | 46 | M | R | 0 | English | Right mid-to-anterior temporal lobe |
| EC250 | L | 30 | M | R | 1 | English | Left medial temporal lobe |
| EC252 | R | 36 | M | R | 0 | Spanish | Right frontoparietal operculum |

**Table S1.** Participant demographic and clinical information.

| **Instrument** | **Source** | **Duration** |
| --- | --- | --- |
| Trumpet | *Stardust*, Hoagy Carmichael/Wynton Marsalis | 0:06 |
| Violin | *Symphony no. 35 in D Major*, W.A. Mozart | 0:07 |
| Saxophone | *Pictures at an Exhibition,* M.P. Mussorgsky | 0:06 |
| Flute | *Orchestral Suite No. 2 in B Minor BWV 1067 -7*, J.S. Bach | 0:09 |
| Bassoon | *Infernal Dance,* Igor Stravinsky | 0:06 |
| Double Bass | *Work in Progress*, Edgar Meyer | 0:05 |
| Cello | *Requiem Offertorio*, Guiseppe Verdi | 0:07 |
| Guitar | *Asturias*, Isaac Albeniz | 0:06 |
| French Horn | *Concerto no. 4 in E flat Major*, W.A. Mozart | 0:04 |
| Electric Bass | *Improvisation*, Jaco Pastorius | 0:09 |
| Mandolin | *Mandolin Concerto in C Major*, Antonio Vivaldi | 0:05 |
| Oboe | *Swan Lake suite Op. 20*, P.I. Tchaikovsky | 0:10 |

**Table S2.** Instrumentation, source, and duration for example musical phrases.

**Audio S1.** Example musical phrases that correspond to phrases listed in table S2.

**Audio S2.** Three example speech tokens.

**Audio S3.** Three melodic speech tokens corresponding to the regular speech tokens in AudioS2. Melodic speech tokens are primed by a piano chord to establish the tonality.
